# Wash and the WASH Regulatory Complex function in Nuclear Envelope budding

**DOI:** 10.1101/2019.12.18.881763

**Authors:** Jeffrey M. Verboon, Mitsutoshi Nakamura, Jacob R. Decker, Kerri A. Davidson, Vivek Nandakumar, Susan M. Parkhurst

## Abstract

Nuclear envelope budding is a recently described phenomenon wherein large macromolecular complexes can be packaged inside the nucleus and be extruded through the nuclear membranes, completely bypassing nuclear pores. While factors have been identified both as cargos or actively involved in this process, much remains unknown about the molecules that generate the forces and membrane deformations which appear inherent. Using fluorescence and electron microscopy, biochemical and cell biological assays, and genetic perturbations in the *Drosophila* model, we identify Wash, its regulatory complex, and Arp2/3 as novel players in NE-budding. Surprisingly, Wash’s role in this process is bipotent and, independent of SHRC/Arp2/3, its perturbation disrupts the normal homotypic Lamin A/B meshworks that are necessary for NE-budding to occur. In addition to NE-budding emerging as important in additional cellular processes and organisms, its incredible similarity to herpesvirus egress suggests new avenues for exploration in both normal and disease biology.

## INTRODUCTION

Transport of macromolecules from the nucleus to the cytoplasm is essential for all developmental processes, including the regulation of differentiation, cell proliferation, and aging, and when mis-regulated, is associated with diseases and tumor formation/progression (Burke and Stewart, 2002; Grunwald et al., 2011; Siddiqui and Borden, 2012; Tran et al., 2014). This indispensable process has been thought to occur exclusively through Nuclear Pore Complexes (NPCs), channels that span the double membrane nuclear envelope and provide a critical regulatory step in what exits (and enters) the nucleus (Daneholt, 2001; Grunwald et al., 2011). Recently, Nuclear Envelope (NE-) budding was identified as an alternative pathway for nuclear exit, particularly for large developmentally-required ribonucleoprotein (megaRNP) complexes that would otherwise need to unfold/remodel to fit through the NPCs (Fradkin and Budnik, 2016; Hatch and Hetzer, 2014; Hatch and Hetzer, 2012; Jokhi et al., 2013; Li et al., 2016; Parchure et al., 2017; Speese et al., 2012). In this pathway, large macromolecule complexes, such as megaRNPs, are encircled by the nuclear lamina (type-A and type-B lamins) and inner nuclear membrane, are pinched off from the inner nuclear membrane, cross the perinuclear space, fuse with the outer nuclear membrane, and release the megaRNPs into the cytoplasm (Figure 1A-C). Strikingly, NE-budding shares many features with the nuclear egress mechanism used by herpesviruses, common pathogens that cause and/or contribute to a diverse array of human diseases (Bigalke and Heldwein, 2016; Hagen et al., 2015; Lye et al., 2017; Mettenleiter et al., 2013; Parchure et al., 2017; Roller and Baines, 2017). As viruses often take advantage of pre-existing host pathways for their livelihoods, the parallel between nuclear exit of herpesvirus nucleocapsids and that of megaRNPs suggests that NE-budding may be a general cellular mechanism that elegantly allows for the nuclear export of endogenous megaRNPs and/or other large cargos (cf. (Fradkin and Budnik, 2016; Mettenleiter et al., 2013; Parchure et al., 2017; Roller and Baines, 2017). Indeed, this pathway has also been implicated in the removal of obsolete macromolecular complexes or other material (i.e., large protein aggregates, poly-ubiquitylated proteins) from the nucleus (Jokhi et al., 2013; Ramaswami et al., 2013; Rose and Schlieker, 2012).

**Figure 1.**
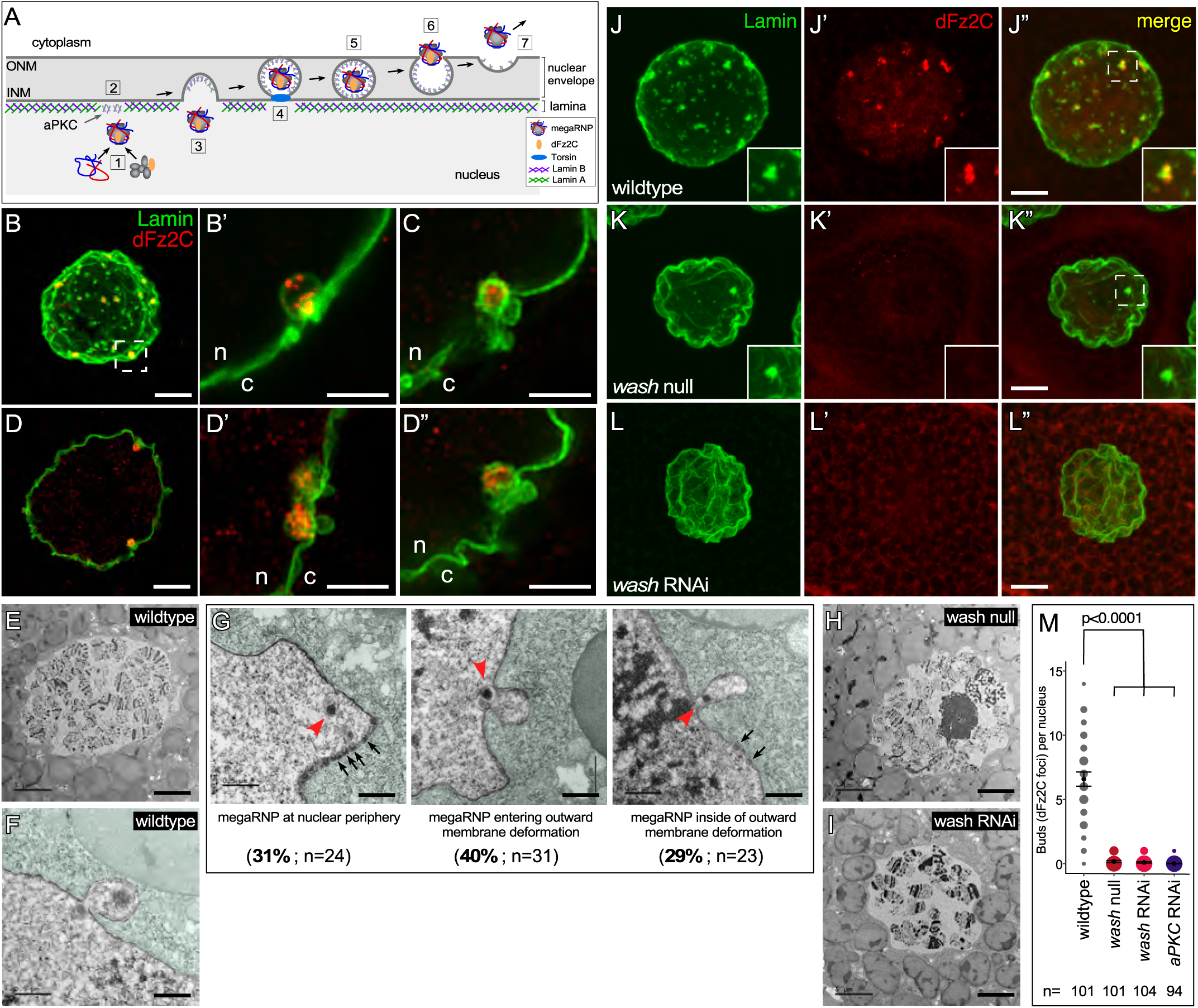
Wash mutant nuclei lack NE-buds. (A) Schematic of NE-budding steps: megaRNPs are assembled and dFz2C is incorporated [1], the nuclear lamina is modified by aPKC [2], megaRNPs enter the membrane deformation [3] and are encapsulated by inner nuclear membrane [4], scission of the inner nuclear membrane [5], inner nuclear membrane fusion with outer nuclear membrane [6], and MegaRNP exit into cytoplasm [7]. (B-D”) Super-resolution micrograph projection (B) or single slice (D) of wildtype larval salivary gland nucleus stained with antibodies to Lamin Band dFz2C. Large Lamin Band dFz2C positive puncta indicate NE-buds (n=nucleus; c=cytoplasm). (B’) High magnification view of highlighted region of (B). (C) High magnification view of NE-buds. (D-D’) High magnification view of NE-buds in (D). (E-F) TEM micrographs at 800X (E) and 8,000X (F) of wildtype larval salivary gland nucleus (cytoplasm false colored in green). (G) TEM micrographs at (8,000X) of NE-buds (red arrowheads) in wildtype larval salivary gland nuclei showing the distribution of phenotypes observed (black arrows denote Nuclear Pore Complexes). (H-I) TEM micrographs (800X) of *wash* null (H) and *wash* RNAi (I) larval salivary gland nuclei. (J-L”) Larval salivary gland nuclei from wildtype (J-J”), *wash* null (K-K”), and *wash* RNAi (L-L”) stained with Lamin B and dFz2C. (M) Quantification of NE-buds per nucleus in larval salivary glands. Kruskal Wallis test; p-values indicated. Scale bars: 5*μ*m in B,D,E-F,H-1,J-L”; 0.5*μ*m in B’-C,D’-D”,G.

The NE-budding pathway was first demonstrated in Drosophila synapse development, proving to be essential for neuromuscular junction integrity. Here, a C-terminal fragment (dFz2C) of the fly Wingless receptor, dFz2C, was shown to associate with megaRNPs that formed foci at the nuclear periphery and exited the nucleus by budding through the nuclear envelope (Figure 1B-C) (Speese et al., 2012). Failure of this process resulted in aberrant synapse differentiation and impaired neuromuscular junction integrity (Speese et al., 2012). In a subsequent study, the NE-budding pathway was shown to be necessary for the nuclear export of megaRNPs containing mitochondrial RNAs such that disruption of NE-budding lead to deterioration of mitochondrial integrity and premature aging phenotypes similar to those associated with A-type lamin mutations (i.e., laminopathies) (Jokhi et al., 2013; Li et al., 2016). Similar endogenous perinuclear foci/buds have been observed in plants and vertebrates, as well as other *Drosophila* tissues, suggesting that cellular NE-budding is a widely-conserved process (cf. (Hadek and Swift, 1962; Hochstrasser and Sedat, 1987; LaMassa et al., 2018; Panagaki et al., 2018; Parchure et al., 2017; Speese et al., 2012; Szollosi and Szollosi, 1988). However, the spectrum of developmental processes requiring this non-canonical nuclear exit pathway and the molecular factors needed for this highly dynamic process, which encompasses membrane deformations, traversal across a membrane bilayer, and nuclear envelope remodeling for a return to homeostasis, remain largely unknown.

Two factors with established roles in NE-budding are aPKC, thought to modify lamins so that budding can occur, and Torsin, needed to pinch off formed buds (Jokhi et al., 2013; Speese et al., 2012). One class of proteins that are involved in membrane-cytoskeletal interactions, regulation, and organization is the Wiskott-Aldrich Syndrome (WAS) protein family (cf. (Takenawa and Suetsugu, 2007). WAS protein subfamilies (WASP, SCAR/WAVE, WASH, WHAMM, JMY) are involved in a wide variety of essential cellular and developmental processes, as well as in pathogen infection, disease, and cancer metastasis (Burianek and Soderling, 2013; Campellone and Welch, 2010; Massaad et al., 2013; Rottner et al., 2010; Rotty et al., 2013; Takenawa and Suetsugu, 2007; Wojnacki et al., 2014). WAS family proteins polymerize branched actin through the Arp2/3 complex, and often function as downstream effectors of Rho family GTPases (Campellone and Welch, 2010; Takenawa and Suetsugu, 2007). We identified WASH as a new WAS subfamily that is regulated in a context-dependent manner: Wash can bind directly to Rho1 GTPase (at least in *Drosophila*) or it can function along with the multi-protein WASH Regulatory Complex (**SHRC**; comprised of SWIP, Strumpellin, FAM21, and CCDC53) (Derivery et al., 2009; Duleh and Welch, 2010; Gomez and Billadeau, 2009; Jia et al., 2010; Linardopoulou et al., 2007; Liu et al., 2009; Park et al., 2013; Veltman and Insall, 2010; Verboon et al., 2018; Verboon et al., 2015a; Verboon et al., 2015b). Wash regulation by Rho family GTPases outside of *Drosophila* has not yet been described (Jia et al., 2010), instead its regulation has only been characterized in the context of its SHRC. WASH and its SHRC are evolutionarily conserved and their mis-regulation is linked to cancers and neurodegenerative disorders (Leirdal et al., 2004; Linardopoulou et al., 2007; McGough et al., 2014; Nordgard et al., 2008; Ropers et al., 2011; Ryder et al., 2013; Turk et al., 2017; Valdmanis et al., 2007; Vardarajan et al., 2012; Zavodszky et al., 2014a; Zavodszky et al., 2014b). Importantly, we have shown that WASH is present in the nucleus where it affects global nuclear organization/functions, interacts directly with B-type lamins, and when mutant exhibits disrupted subnuclear structures/organelles, as well as abnormal wrinkled nucleus morphology reminiscent of that observed in diverse laminopathies (Verboon et al., 2015b; Verboon et al., 2015c). Mammalian WASH proteins have also been shown to localize to the nucleus in developmental and cell-type specific manners (Verboon et al., 2015c; Xia et al., 2014).

Here we show that WASH and its SHRC are also involved in the NE-budding pathway, as *wash* and SHRC mutants lack dFz2C foci/lamin buds and display the neuromuscular junction integrity and premature aging phenotypes previously associated with the loss of NE-budding. In addition, we find that CCDC53 (SHRC subunit) co-localizes with dFz2C foci/lamin buds. We show that Wash is present in several independent nuclear complexes. Wash’s nuclear interactions with its SHRC are separate of those with B-type Lamin, leading to effects on different subsets of nuclear Wash functions. We also find that WASH-dependent Arp2/3 actin nucleation activity is required for proper NE-budding. We propose that Wash and its SHRC play a physical and/or regulatory role in the process of NE-budding.

## RESULTS

### Wash mutants lack NE-buds

*Drosophila* larval salivary glands undergo NE-budding: dFz2C-containing megaRNPs can be observed surrounded by Lamin B at the nuclear periphery using confocal microscopy (Figure 1B-D”, S1A) or Transmission Electron Microscopy (TEM; Figure 1E-F) (Hochstrasser and Sedat, 1987). Using TEM, we observe that these megaRNPs are adjacent to a curved evagination of the nuclear membrane, suggesting that there is likely a membrane deformation event that precedes megaRNP encapsulation by the inner-nuclear envelope (40%, n=78 NE-budding events) (Figure 1G). Finally, dFz2C antibody co-localizes in a subset of these Lamin foci suggesting that dFz2C can act as a biomarker for nuclear buds in the salivary gland (Figure 1B-D”, J-J”).

While staining *wash* mutants for Lamin B previously, we observed notably fewer Lamin ‘buds’ (for either A- or B-type Lamins) than in wildtype (Verboon et al., 2015b). To determine if this reduction of Lamin buds was a result of disrupted NE-budding, we co-labeled *wash* mutant larval salivary gland nuclei for Lamin B and dFz2C. We generated *wash* mutant salivary glands two different ways: 1) using an outcrossed homozygous null allele, *wash^Δ185hz(outX)^* (hereafter referred to as “*wash* null”; (Verboon et al., 2018), and 2) expressing an RNAi construct for *wash* (HMC05339) specifically in the salivary gland using the GAL4-UAS system (hereafter referred to as “*wash* RNAi”) (Figure 1J-L”). We find that *wash* null and *wash* RNAi larval salivary gland nuclei exhibit an average of 0.2±0.0 (n=101) and 0.1±0.0 (n= 104) dFz2C foci/NE-buds respectively, compared to an average of 6.6±0.3 dFz2C foci/NE-buds in wildtype (n=101, p<0.0001) (Figure 1J-L”). The phenotypes that we observe are similar to those reported previously for aPKC knockdowns (0.0±0.0, n=94, Figure 1M) (Fradkin and Budnik, 2016; Jokhi et al., 2013; Li et al., 2016; Parchure et al., 2017). TEM sections from *wash* null and *wash* RNAi knockdown nuclei showed a wrinkled nuclear membrane and a similar lack of NE-buds (n=50, n=50 respectively; Figure 1H-I).

### Wash mutants display phenotypes associated with loss of NE-budding

As our results indicate that Wash is involved in NE-budding in the salivary gland, and if Wash is a conserved factor in this process, we would expect *wash* mutants to exhibit the phenotypes associated with the two processes shown previously to be reliant on NE-budding: neuromuscular junction development and mitochondrial integrity.

Motor neurons in the body larval wall muscle have synaptic boutons, which are glutamatergic and are necessary for contractility of this muscle (Menon et al., 2013). In wildtype larvae these boutons are largely mature and contain only on average 1.6±0.1 per 100 boutons (n=664) exhibiting an absence of the neuronal differentiation marker *disks large* (DLG) or “ghost bouton” phenotype (Figure 2A, D). Motor neurons from *wash* null and *wash* RNAi larval body wall muscle have an increased number of ghost boutons: 8.5±0.9 ghost boutons per 100 boutons (n=570, p=<0.0001) and 8.4±0.5 ghost boutons per 100 boutons (n=649, p=<0.0001), respectively (Figure 2B-D), consistent with phenotypes observed under conditions known to disrupt nuclear budding (Speese et al., 2012).

**Figure 2.**
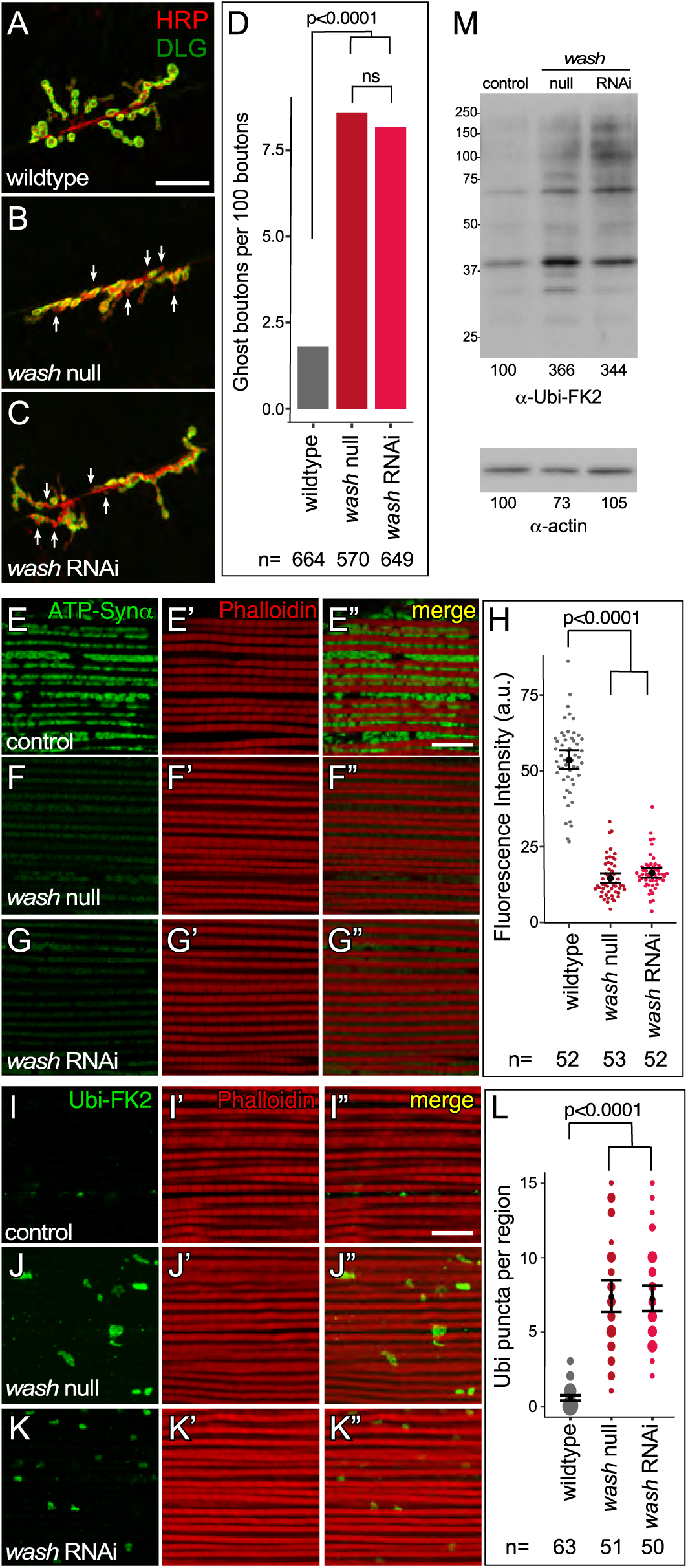
Wash mutant nuclei display NE-bud associated phenotypes. (A-C) Confocal micrograph projection of synaptic boutons from wildtype (A), *wash* null (B), and *wash* RNAi (C) larval body wall muscle showing HRP and DLG expression. Synaptic boutons from *wash* null and *wash* RNAi larvae display ghost bouton phenotype (loss of DLG staining; arrows). (D) Quantification of ghost bouton frequency in larval body wall muscle neurons. (E-G”) Confocal micrograph projection of adult indirect flight muscle (IFM) from wildtype (E-E”), *wash* null (F-F”),and *wash* RNAi (G-G”) flies aged 21 days stained with activity dependent mitochondrial marker ATP-Syn α and Phalloidin. (H) Quantification of mitochondrial fluorescence intensity in IFM. (I-K”) Confocal micrograph projection of adult IFM from wildtype (I-I”), *wash* null (J-J”), and *wash* RNAi (K-K”) flies aged 21 days stained with poly-ubiquitin aggregate marker Ubi-FK2 and Phalloidin. (L) Quantification of number of large Ubi-FK2 marks per muscle region in IFM. (M) Western blot of adult IFM lysates from control, *wash* null, and *wash* RNAi flies aged 21 days showing poly-ubiquitin aggregate protein levels, and actin loading control. Two-tailed Fisher’s exact test (D); Two-tailed student’s t-test (H); Kruskal Wallis test (L); all p-values indicated. Scale bars: 20μm in A-C; 10*μ*m in E-G”,1-K”.

We next looked at whether *wash* affects mitochondrial integrity in the indirect flight muscle (IFM) associated with the NE-budding process using ATP-Syn α. *wash* null and *wash* RNAi IFMs from adults aged 21 days show a 3.7 (n=53) and 3.2 (n=52) fold decrease respectively (p<0.0001), in mitochondrial activity compared to a wildtype control (Figure 2E-H). IFM from *wash* null and *wash* RNAi flies aged 21 days also show an increase in poly-ubiquitin aggregates (a marker of mitochondrial damage), as muscle regions from these mutants show an average of 7.7 (n=51) and 7.4 (n=50) large poly-ubiquitin positive puncta, while IFM from control flies show just 0.5 (n=63) large poly-ubiquitin positive puncta on average (Figure 2I-L). Tissue lysates from *wash* null and *wash* RNAi IFMs from adults aged 21 days show a similar increase of poly-ubiquitin aggregates by western blot, indicating mitochondrial damage (Figure 2M). Taken together these data indicate that Wash plays a fundamental role in NE-budding.

### SHRC regulates Wash activity in NE-budding and accumulates at NE-budding sites

*Drosophila* Wash undergoes context-dependent regulation: it can function as a direct effector of the Rho1 small GTPase, or it can function as part of the SHRC, while a pathway where Wash interacts with both has not been demonstrated (Liu et al., 2009; Verboon et al., 2018; Verboon et al., 2015a). We had previously shown that Wash localizes to both the cytoplasm and nucleus in salivary glands (Verboon et al., 2015b). To determine if Rho1 or SHRC were working with Wash in the context of NE-budding, we stained larval salivary glands with antibodies specific to Wash, Rho1, and each of the four SHRC components (CCDC53, Strumpellin, SWIP, and FAM21) (Magie et al., 2002; Rodriguez-Mesa et al., 2012; Verboon et al., 2015a). We find that, as expected, Wash, Rho1, and all 4 SHRC proteins are expressed throughout the cytoplasm and enriched at cell boundaries (Figure 3A-F”). Interestingly, while Wash and all 4 SHRC proteins are expressed highly in the nucleus, Rho1 is not (Figure 3A-F”, S1B), indicating that the SHRC could regulate Wash activity in the context of NE-budding. Importantly, the SHRC component CCDC53 co-localizes with dFz2C at NE-buds (Figure 3G-H”). CCDC53 also co-localizes with the Lamin B that encapsulates NE-buds (Figure 3I-K”) and is particularly enriched at the base (neck) of NE-buds suggesting a direct nuclear role for these proteins (Figure 3H-H”, K-K”). Notably, we do not observe enrichment of the other SHRC subunits at the nuclear periphery coincident with NE-buds. It could be that the epitopes recognized by the other SHRC component antibodies are masked at these sites, or it could represent an activity of CCDC53 that is independent of Wash and its SHRC, however we favor the former given the remarkable consistency between nuclear budding phenotypes of CCDC53 and Wash.

**Figure 3.**
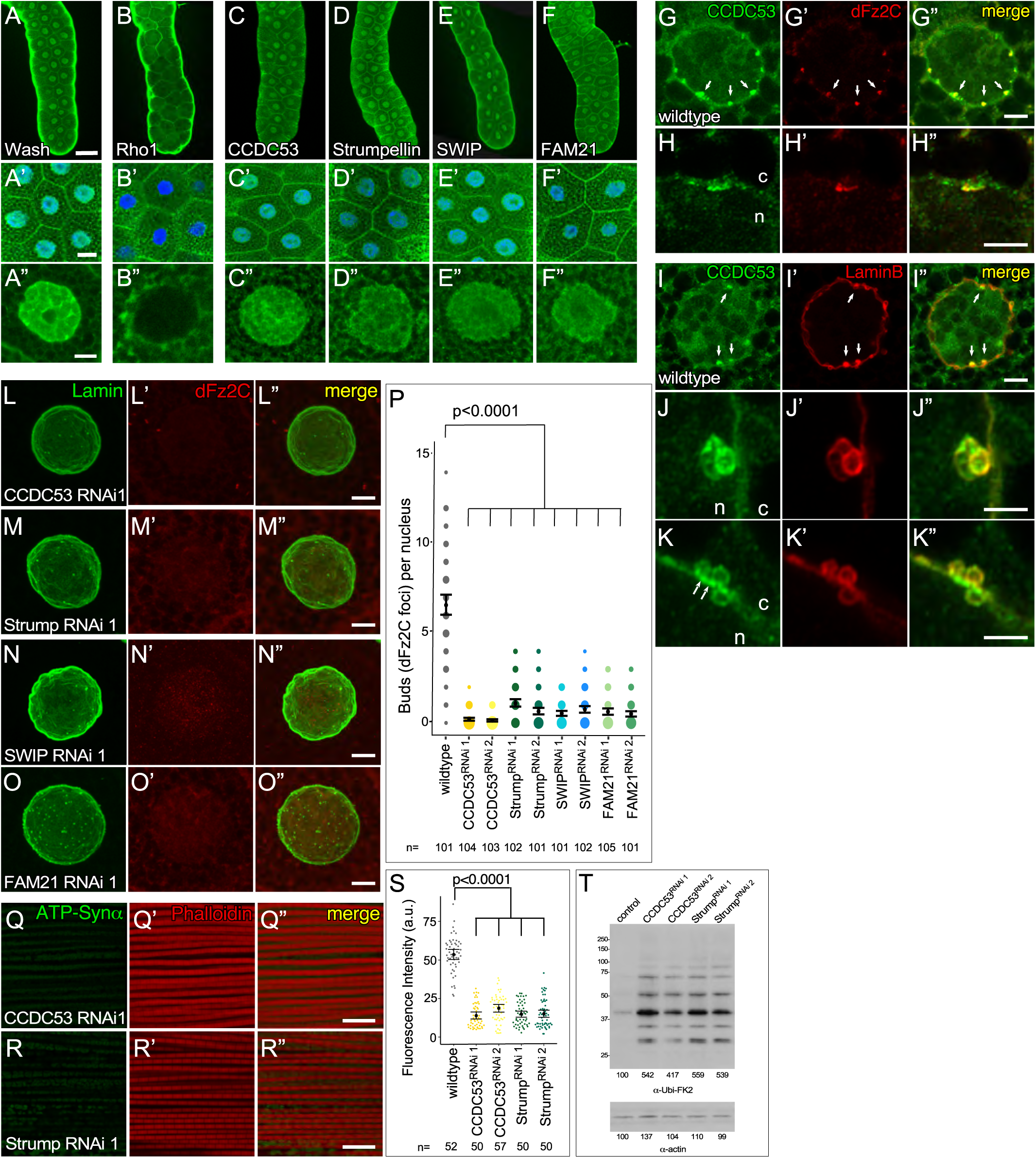
SHRC is expressed in the nucleus, accumulates at NE-buds, and SHRC mutants lack NE-buds and display downstream NE-budding associated phenotypes. (A-F”) Confocal micrograph projection of wildtype larval salivary glands showing expression of Wash (A-A”), Rho1 (B-B”), and SHRC subunits: CCDC53 (C-C”), Strumpellin (D-D”), SWIP (E-E”), and FAM21 (F-F”) at cell boundaries, throughout cytoplasm and in nuclei. DAPI staining (blue) marks nuclei in (B’-F’). (G-G”) Single slice super-resolution (Airyscan) micrograph of wildtype larval salivary gland nucleus showing CCDC53 enrichment at NE-bud sites and colocalization with dFz2c (arrows). (H-H”) High magnification view of single budding site from (G). (I-K”) Single slice super-resolution micrograph of wildtype larval salivary gland nucleus showing that CCDC53 is enriched at NE-bud sites and colocalizes with Lamin B puncta at the nuclear periphery (arrows). (J-K”) High magnification views of CCDC53 colocalization with Lamin B at NE-buds from (I). (L-O”) Confocal micrograph projections of CCDC53 RNAi (L-L”), Strumpellin RNAi (M-M”), SWIP RNAi (N-N”), and FAM21 RNAi (O-O”) larval salivary gland nuclei stained with Lamin B and dFz2C. (P) Quantification of NE-buds per nucleus from two independent RNAi lines per SHRC subunit. (Q-R”) Confocal micrograph projections of adult IFM from CCDC53 RNAi (Q-Q”) and Strumpellin (R-R”) RNAi flies aged 21 days stained with activity dependent mitochondrial marker ATP-Syn α and Phalloidin. (S) Quantification of ATP-Syn α fluorescence intensity in adult IFM. (T) Western blot of adult IFM lysates from control, CCDC53 RNAi, and Strumpellin RNAi flies aged 21 days showing poly-ubiquitin aggregate protein levels, and actin loading control. Two-tailed student’s t-test (S); Kruskal Wallis test (P); all p-values indicated. Scale bars: 100μm in A-F; 20μm in A’-F’; 5μm in A”-G”,I-I”,L-O”; 0.5μm in H-H”,J-K”; 10μm in Q-R” (n=nucleus; c=cytoplasm).

To determine if SHRC activity is required for NE-budding, we knocked down individual SHRC components by expressing RNAi constructs (two independent RNAi constructs for each subunit) specifically in the larval salivary gland using the GAL4-UAS system (Figure 3L-O”; Figure S1C-F”). We co-labeled these SHRC knockdown larval salivary gland nuclei for Lamin B and dFz2C and find that all four SHRC knockdowns show a significant decrease in NE-buds similar to that of *wash* null and *wash* RNAi larval salivary gland nuclei (Figure 3L-O”; Figure S1C-F”). The number of dFz2C foci/NE-buds per nucleus was quantified for each line (Figure 3P) and we find that all lines show 0.1-1.1±0.1 foci/buds per nucleus on average (n>100 and p<0.0001 in each case) compared to 6.6±0.3 foci/buds in control nuclei (n=100, p<0.0001). Of note, the nuclei from SHRC knockdown larval salivary glands are spherical and morphologically indistinct from wildtype nuclei, suggesting that the wrinkled nuclear phenotype observed in *wash* mutants (Verboon et al., 2015b) is SHRC independent.

Consistent with their loss of dFz2C foci/NE-buds, SHRC component knockdowns also exhibit phenotypes associated with disrupted NE-budding (Figure 3Q-T). Adult IFM from 21 day old *Strump* or *CCDC53* knockdown flies show a decrease in mitochondrial activity, measured by the activity-dependent mitochondrial marker ATP-Synthetase α (Figure 3Q-S; Figure S1G-H”). *Strump* RNAi1 and RNAi2 show a 3.6 and 3.5 fold decrease in activity compared to wildtype (n=50 and n=50, p<0.0001) and *CCDC53* RNAi1 and RNAi2 show a 3.8 and 2.9 fold decrease in activity compared to wildtype (n=50 and n=57, p<0.0001) (Figure 3S). Western blots of IFM lysates from these knockdowns also show an increase in poly-ubiquitin aggregates, indicating mitochondrial damage (Figure 3T). Taken together, our results are consistent with nuclear Wash functioning with its SHRC in NE-budding.

### Lamin meshes are required for nuclear budding

We were next interested in understanding how Wash and its regulation by the SHRC were involved in the dynamic process of NE-budding. As it has been previously reported that mutations in Lamin C (A-type) disrupt NE-budding (Li et al., 2016; Parchure et al., 2017), and since we had shown that Wash interacts directly with Lamin B (B-type) (Verboon et al., 2015b), we wanted to determine whether or not the defect in NE-budding was due to a structural defect in Lamin B organization via its interaction with Wash. Lamin B and Lamin A/C have been shown to form homotypic meshes underlying the inner nuclear membrane (Shimi et al., 2015) and, due to their intimate relationship, might be expected to affect NE-budding. Indeed, salivary gland nuclei from *Lamin B* RNAi knockdowns exhibit a characteristic wrinkled nuclear morphology and a decrease in dFz2C foci/NE-buds, with 2.2±0.1 buds per nuclei (n=171) (Figure 4A-B). Consistent with this reduction in NE-buds, adult IFMs from *Lamin B* RNAi flies aged 21 days show an 3.1 fold decrease in mitochondrial activity from wildtype (n=50, p<0.0001), as assayed by the activity dependent mitochondrial marker ATP-Synthetase α (Figure 4C-D). Additionally, western blots of adult IFM lysates from *Lamin B* RNAi flies aged 21 days show an increase in poly-ubiquitin aggregates, indicating mitochondrial damage (Figure 4E). These data suggest that alteration of nuclear lamina structure can reduce, but not eliminate, NE-budding.

**Figure 4.**
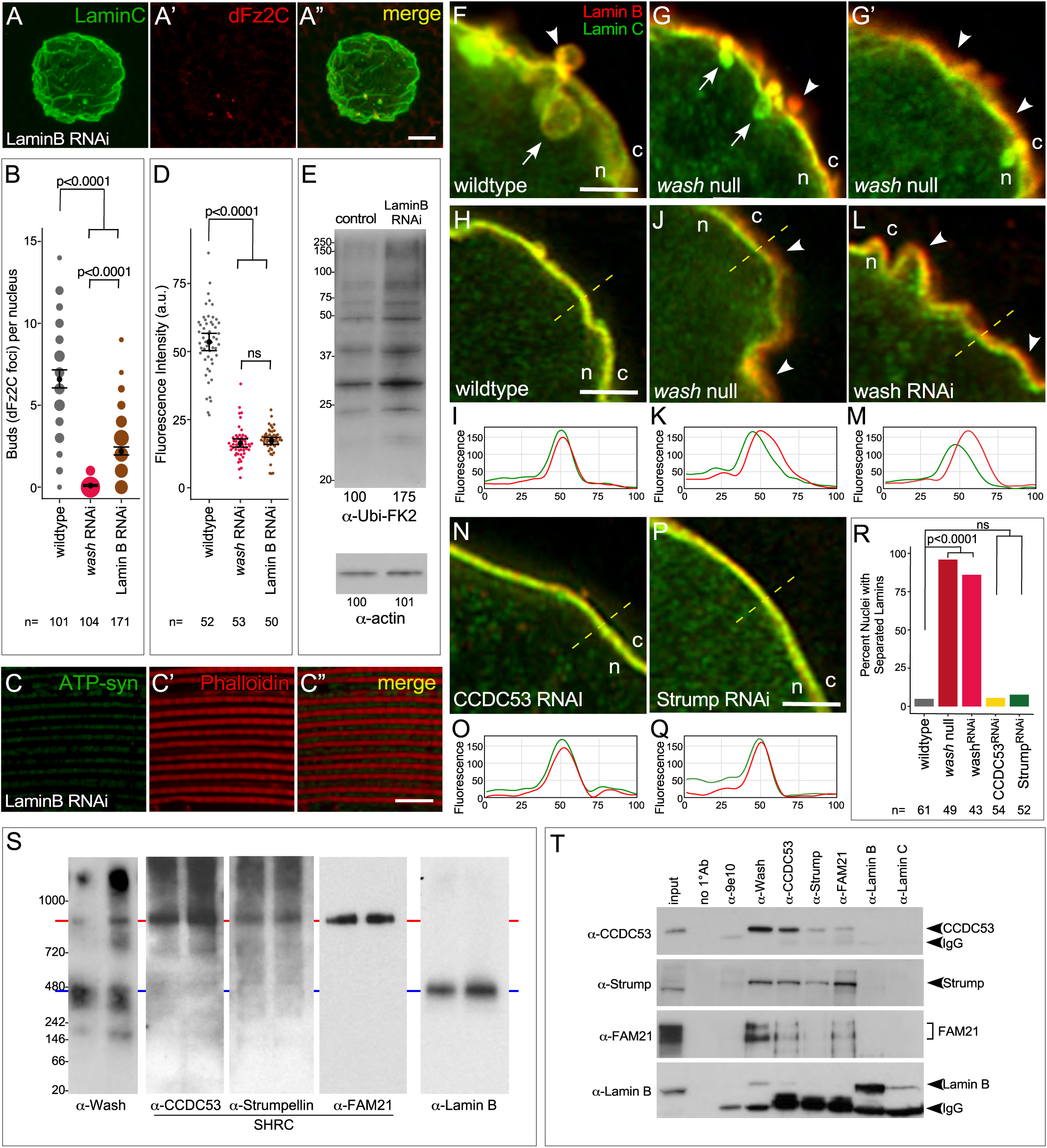
Alteration of the nuclear lamina structure can reduce, but does not eliminate, NE-budding. (A) Confocal micrograph projection of Lamin B RNAi larval salivary gland nucleus co-labeled with antibodies to Lamin C and dFz2C. (B) Quantification of number of NE-buds in control, *wash* RNAi, and Lamin B RNAi nuclei. (C) Confocal micrograph projection of adult IFM from Lamin B RNAi flies aged 21 days stained with ATP-Syn α and Phalloidin. (D) Quantification of ATP-Syn α fluorescence intensity in adult IFM. (E) Western blot of adult IFM lysates from control and Lamin B RNAi flies aged 21 days showing poly-ubiquitin aggregate protein levels, and actin loading control. (F-G) Single slice confocal micrographs of nuclear periphery from wildtype (F) and *wash* null (G-G’). (H-Q) Single slice super-resolution micrographs of larval salivary gland nuclei (H,J,L,N,P) and line plots of region indicated (I,K,M,O,Q) from wildtype (H-I), *wash* null (J-K), *wash* RNAi (L-M), CCDC53 RNAi (N-O), and Strumpellin RNAi (P-Q) showing Lamin B/Lamin C organization at the nuclear periphery. Lamin C also shows lower level uniform distribution within the nucleus. (R) Quantification of the percentage of salivary gland nuclei showing Lamin B/Lamin C separation. (S) Western blots from Blue Native PAGE of nuclear extracts probed with antibodies to Wash, SHRC subunits (CCDC53, Strumpellin, and FAM21), and Lamin B. Putative ∼900KDa complex with Wash and SHRC (red line) and ∼450KDa complex with Wash and Lamin B (blue lines) are indicated. (T) Western blots of immunoprecipitations from nuclear extracts with no primary antibody included (no 1° AB), a non-specific antibody (9e10), Wash, CCDC53, Strumpellin, FAM21, Lamin B, and Lamin C. Blots were probed with antibodies to CCDC53, Strumpellin, FAM21, and Lamin B as indicated. Two-tailed Fisher’s exact test (R); Two-tailed student’s t-test (D); Kruskal Wallis test (B); all p-values indicated. Scale bars: 5μm in A-A”; 1μm in F-H,J,L,N,P; 10μm in C-C” (n=nucleus; c=cytoplasm).

### Wash, but not the SHRC, interacts with and disrupts homotypic Lamin meshes

We were next interested in exploring whether or not *wash* and/or *SHRC* genotypes caused budding defects by causing defects in Lamin organization. Interestingly, our previous data suggested that Wash’s involvement with Lamin B might be disparate from its involvement with the SHRC, as Wash mutants exhibit a wrinkled nuclear morphology (Verboon et al., 2015b), whereas SHRC mutants do not (Figure 1K-L’’ versus Figure 3L-O’’). Thus, we analyzed the structure of the nuclear lamina in these different mutants. As previously mentioned, Lamin B and Lamin A/C have been shown to form homotypic meshes, but how these different lamin networks are organized with respect to each other, as well as the inner nuclear membrane, is not well understood (Shimi et al., 2015). In particular, factors that disrupt this homotypic meshwork may disrupt the structural platform at/upon which NE-budding occurs. Wildtype larval salivary gland nuclei co-labeled with antibodies to Lamin B and Lamin C show tight Lamin B and Lamin C colocalization around the nuclear periphery (Figure 4F,H-I,R). However, in nuclei from *wash* null and *wash* RNAi larval salivary glands, the signals from the two Lamins are separated with Lamin B lying closest to the inner nuclear membrane (consistent with it encoding a CAAX domain) and Lamin C positioned closest to the chromatin (consistent with Lamin C association with chromatin) (Figure 4G-G’,J-M,R). Importantly, nuclei from SHRC subunit knockdowns, *CCDC53* RNAi and *Strump* RNAi, have Lamin B/Lamin C organization that is indistinct from wildtype (Figure 4N-R).

As an orthogonal means of determining if Wash functions separately of the SHRC to influence the nuclear lamina, we separated protein complexes from fly cell nuclear lysates using blue native-PAGE. We observe that Wash is present in multiple nuclear complexes, suggesting it is involved in multiple nuclear processes (Figure 4S). We find that one major Wash-containing complex (∼900KDa) overlaps with a complex containing CCDC53, Strumpellin, and FAM21 (Figure 4S), whereas a separate Wash-containing complex (∼450 KDa) overlaps with a Lamin B-containing complex (Figure 4S). To confirm that Wash forms distinct complexes with the SHRC components and with Lamin B in the nucleus, we immunoprecipitated Wash, three SHRC subunits (CCDC53, Strumpellin, and FAM21), Lamin B, and Lamin C from fly cell nuclear lysates and probed the resulting western blots for SHRC and Lamin B (Figure 4T). In each case, Wash pulls down the corresponding SHRC subunit, the SHRC subunit pulls itself down, and SHRC complex members co-immunoprecipitated with each other (Figure 4T). However, while Wash pulls down Lamin B, none of the SHRC subunits co-immunoprecipitated with Lamin B or Lamin C. These results are consistent with Wash forming distinct nuclear complexes with Lamin B and with the SHRC. Taken together, our results suggest that Wash may have two roles in NE-budding: an indirect role owing to regulating the Lamin meshwork through its interaction with Lamin B, and a separate role wherein Wash and its SHRC may be directly involved in the physical formation of NE-buds.

### Wash-SHRC interactions are required for NE-budding, whereas Wash-Lamin B interaction is required for nuclear morphology

To determine whether we could separate Wash’s function in NE-budding via interaction with Lamin B versus with its SHRC, we mapped the sites on the Wash protein that facilitate binding to CCDC53 (Wash’s interaction with this SHRC subunit has been reported to facilitate stable Wash-SHRC complex formation; (Derivery et al., 2009; Jia et al., 2010; Rottner et al., 2010; Verboon et al., 2018) and Lamin B, then made specific point mutations that functionally block these interactions (see Methods) (Figure S1I). The final constructs, harboring point mutations in the context of the full-length Wash protein, designated wash^ΔSHRC^ and wash^ΔΔLamB^, respectively, were examined for interaction specificity using GST pulldown assays (Figure S1J). Transgenics generated with these Wash point mutations (see Methods), as well as a wildtype Wash rescue construct (*wash^WT^*), were individually crossed into the *wash* null homozygous background so that the only Wash activity comes from the transgene under control of the endogenous *wash* promoter. The wildtype version of these transgenics (*wash^WT^*) is expressed in both the nucleus and cytoplasm (Verboon et al., 2015b) and rescues previously described *wash* mutant phenotypes, including its premature ooplasmic streaming phenotype in oocytes (Figure S1K-K’, O) (Liu et al., 2009; Verboon et al., 2018). The other two *wash* point mutation transgenic lines are similarly functional: as expected, *wash^ΔΔLamB^* rescues the premature ooplasmic streaming phenotype, whereas *wash^ΔSHRC^* does not (Figure S1L-M’, O).

Larval salivary gland nuclei from the *wash^WT^* show a rescue of NE-budding: nuclei from these mutants show 6.6±0.3 buds per nuclei (n=102; p=0.895 compared to wildtype) and nuclear morphology is indistinct from wildtype (Figure 5A-A”,D). However, nuclei from *wash^ΔSHRC^* point mutants show 0.5±0.1 buds per nucleus (n=101, p<0.0001), but had nuclear morphology indistinct from *wash^WT^* (Figure 5C-D). These data strongly suggest that Wash, under regulatory control of the SHRC, is required for NE-budding. Remarkably, larval salivary gland nuclei from the *wash^ΔΔLamB^* point mutants (Figure 5B-B”, D) exhibit an intermediate phenotype with 1.5±0.1 NE-buds per nucleus, a significantly greater number of NE-buds than that exhibited by the *wash^ΔSHRC^* point mutant (n=104, p<0.0001). *wash^ΔΔLamB^* also exhibit a wrinkled nuclear morphology similar to *wash* null mutants, *wash* RNAi knockdowns, and *Lamin B* RNAi knockdowns (Figure 5B) (Verboon et al., 2015b). Super-resolution microscopy analysis of Lamin B and Lamin C organization in salivary gland nuclei from the *wash* point mutants shows that *wash^ΔΔLamB^* exhibits separated Lamin B and Lamin C meshes, while *wash^WT^* and *wash^ΔSHRC^* have Lamin organization indistinct from wildtype consistent with the specificity of these mutations (Figure 5E-K).

**Figure 5.**
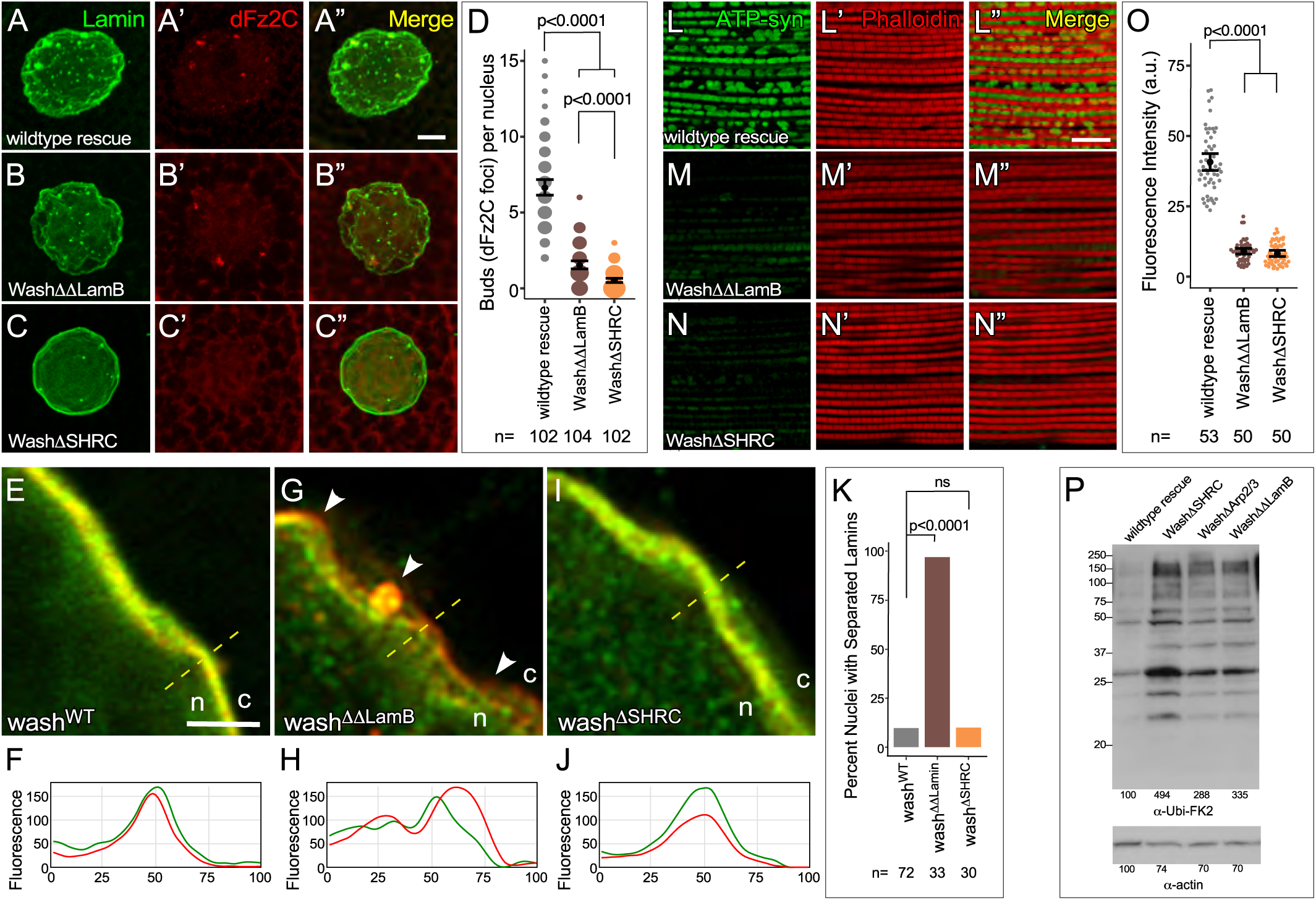
Wash point mutants show separation of phenotypes for specific Wash activities. (A-C”) Confocal micrograph projections of larval salivary gland nuclei from *wash*^WT^ (A-A”), *wash*^ΔΔLamB^ (B-B”), and *wash*^ΔSHRC^ (C-C”) stained with Lamin Band dFz2C. (D) Quantification of NE-buds per nucleus in larval salivary gland nuclei. (E-J) Single slice super-resolution micrographs of larval salivary gland nuclei (E,G,I) and line plots of region indicated (F,H,J) from *wash*^WT^ (E-F), *wash*^ΔΔLamB^ (G-H), and *wash*^ΔSHRC^ (I-J) showing Lamin B/Lamin C organization at the nuclear periphery (n=nucleus; c=cytoplasm). (K) Quantification of the percentage of salivary gland nuclei showing Lamin B/Lamin C separation. (L-N”) Confocal micrograph projections of adult IFM from *wash*^WT^ (L-L”), *wash*^ΔΔLamB^ (M-M”), and *wash*^ΔSHRC^ (N-N”) flies aged 21 days stained with activity dependent mitochondrial marker ATP-Syn α and Phalloidin. (O) Quantification of ATP-Syn α fluorescence intensity from adult IFMs. (P) Western blot of adult IFM lysates from *wash*^WT^, *wash*^ΔΔLamB^, *wash*^ΔSHRC^ and *wash*^ΔArp2/3^ flies aged 21 days showing poly-ubiquitin aggregate protein levels, and actin loading control. Two-tailed Fisher’s exact test (K); Two-tailed student’s t-test (O); Kruskal Wallis test (D); all p-values indicated. Scale bars: 5*μ*m in A-C”; 1*μ*m in E,G,I; 10*μ*m in L-N” (n=nucleus; c=cytoplasm).

As expected due to an overall reduction in NE-buds, IFM from 21 day old *wash^ΔSHRC^* and *wash^ΔΔLamB^* point mutants both show a decrease in mitochondrial activity, as assayed using the activity dependent mitochondrial marker ATP-Synthetase α (Figure 5L-O). IFM’s from *wash^ΔSHRC^* and *wash^ΔΔLamB^* show a 5.0 (n=50) and 4.5 (n=50) fold decrease in mitochondrial activity, respectively, when compared to IFMs from the *wash^WT^* construct (p<0.0001 in each case). Additionally, western blots of IFM lysates from *wash^ΔSHRC^* and *wash^ΔΔLamB^* show an increase in poly-ubiquitin aggregates compared to the *wash^WT^* construct, indicating mitochondrial damage (Figure 5P). Taken together, these point mutations clearly demonstrate Wash’s dual roles in NE-budding as a required element for proper organization of the nuclear lamina compartment where budding occurs and through a disparate mechanism involving its SHRC.

### Wash is required for the initial steps of NE-bud formation

We were next interested in understanding how Wash and its SHRC fits into the physical budding pathway by its relation to known cellular events comprising NE-budding. It has previously been shown that aPKC is necessary to phosphorylate Lamin and seed sites that can undergo NE-budding (Speese et al., 2012). As both *wash* and *aPKC* almost completely abolish NE-budding, epistasis experiments are not feasible. In addition, knockdown of Wash downregulates expression of all four SHRC members (Verboon et al., 2018). However, because the SHRC member CCDC53 accumulates at budding sites, we looked at localization of CCDC53 in aPKC knockdown salivary glands. Consistent with aPKC’s role allowing for this process to occur, CCDC53 does not accumulate in foci nor are NE-buds present in this background (Figure 6A-A”).

**Figure 6.**
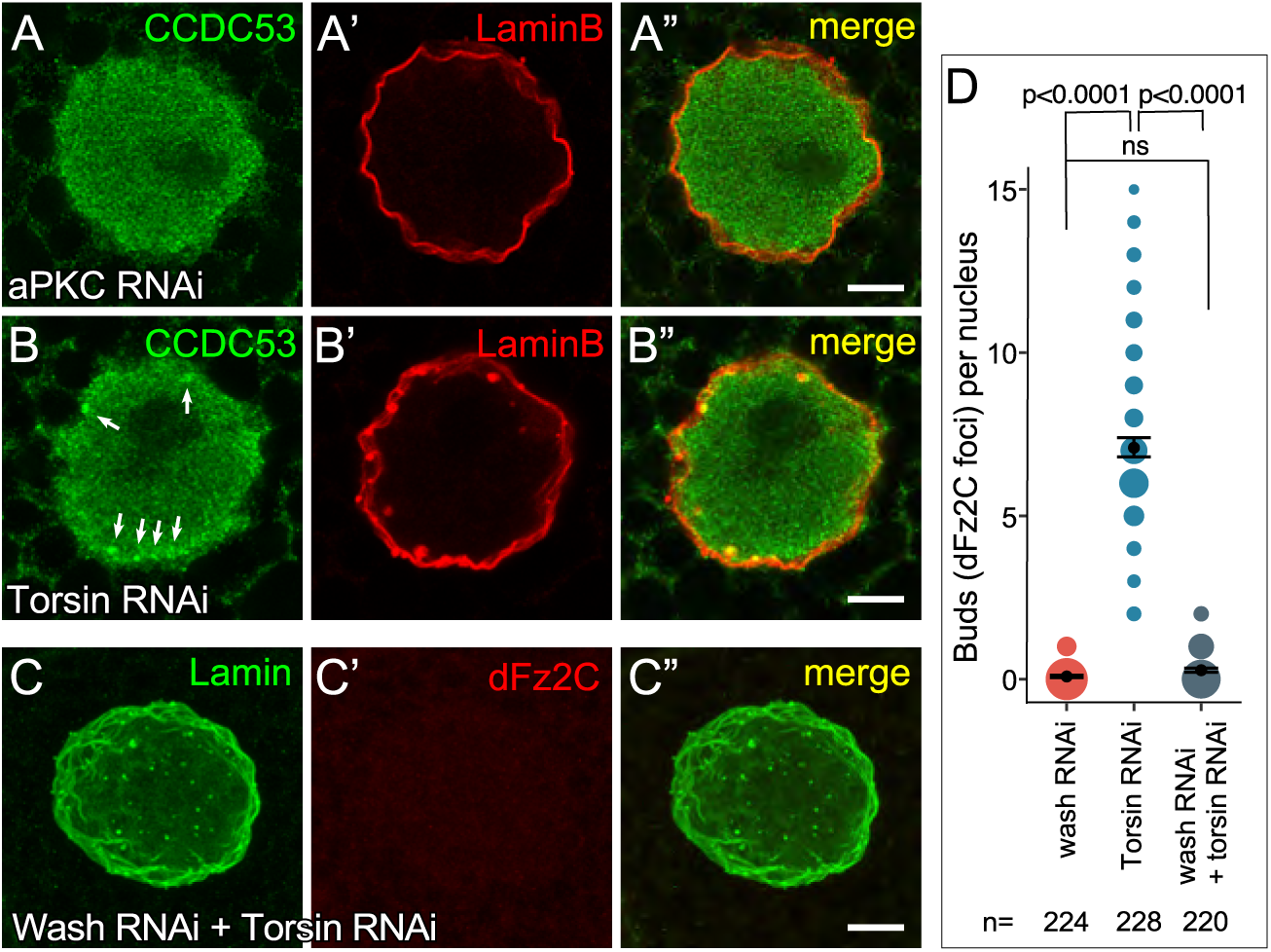
Wash acts prior to NE-buds being pinched off from the inner nuclear membrane. (A-A”) Single slice super-resolution micrograph from an aPKC RNAi larval salivary gland nucleus showing loss of CCDC53 enrichment at NE-bud sites or colocalization with Lamin Bat the nuclear periphery. (B-B”) Single slice super-resolution micrograph from a Torsin RNAi larval salivary gland nucleus showing CCDC53 enrichment at NE-bud sites and colocalization with Lamin B at the nuclear periphery. (C-C”) Confocal micrograph projection of larval salivary gland nuclei from a Wash RNAi +Torsin RNAi double knockdown stained with Lamin Band dFz2C. (D) Quantification of number of dFz2C foci/NE-buds in *wash* RNAi, *torsin* RNAi, and *wash RNAi+torsin* RNAi nuclei. Kruskal Wallis test; p-values indicated. Scale bars: 5*μ*m.

Torsin, an AAATPase, has been proposed to function in the pinching off of the NE-buds from the INM (Jokhi et al., 2013). To determine if Wash and its SHRC is required for the initial steps of NE-bud formation and in particular before these buds pinch off from the INM, we also looked at the localization of CCDC53 in *torsin* RNAi knockdown salivary gland nuclei and find increased numbers of NE-buds co-localizing with CCDC53 foci (Figure 6B-B”), suggesting that torsin acts downstream of Wash’s role in this pathway. To verify this, we generated *wash* and *torsin* double RNAi knockdown larval salivary gland nuclei and co-stained them for Lamin B and dFz2C. We find that *wash* and *torsin* double RNAi larval salivary gland nuclei exhibit an average of 0.3±0.0 dFz2C foci/NE-buds (n=220), compared to an average of 6.6±0.3 dFz2C foci/NE-buds in wildtype (n=100, p<0.0001) (Figure 6C-D) and 0.1±0.0 and 7.1±0.2 dFz2C foci/NE-buds in *wash* RNAi and *torsin* RNAi, respectively. Taken together, these results demonstrate that Wash and the SHRC act in the early steps of NE-budding, between modification of the Lamin at NE-budding sites and the buds pinching off from the INM.

### The Arp2/3 complex is required for NE-budding

As WAS family proteins have been implicated in membrane-cortical cytoskeleton interactions leading to membrane deformations (i.e., protrusions, endocytosis) in the cytoplasm, we explored whether or not it might have a similar role in NE-budding. Wash encodes several biochemical activities, including actin nucleation and actin/MT binding, bundling, and crosslinking (Liu et al., 2009; Verboon et al., 2018; Verboon et al., 2015a); however, when working in concert with the SHRC, Wash often functions upstream of the Arp2/3 complex to promote branched actin filament formation (cf. (Alekhina et al., 2017; Rottner et al., 2010). Mounting evidence has also demonstrated roles for actin and Arp2/3 complexes in diverse nuclear functions (Alekhina et al., 2017; Kyheroinen and Vartiainen, 2019; Percipalle and Vartiainen, 2019). If Wash requires its actin nucleation activity for NE-budding, we would expect Arp2/3 complex subunit mutations to exhibit reduced NE-bud formation and phenotypes associated with disrupted NE-budding. We examined the expression and knockdown effects of two different Arp2/3 complex subunits. The Arp3 and Arpc1 subunits are expressed in fly cell nuclear lysates as assayed by western blot (Figure 7A-B). We generated RNAi knockdowns of *Arp3* and *Arpc1* in larval salivary glands as previously described and find that these nuclei had on average 0.8±0.1 (n=102) and 1.0±0.1 buds (n=102), respectively, compared to 6.6±0.3 buds per nucleus in wildtype (n=102; p<0.0001 in each case) (Figure 7C-E). Additionally, nuclei from *Arp3* and *Arpc1* knockdown animals are spherical (similar to wildtype and SHRC knockdowns), consistent with Wash-dependent Arp2/3 activity being separate from nuclear morphology.

**Figure 7.**
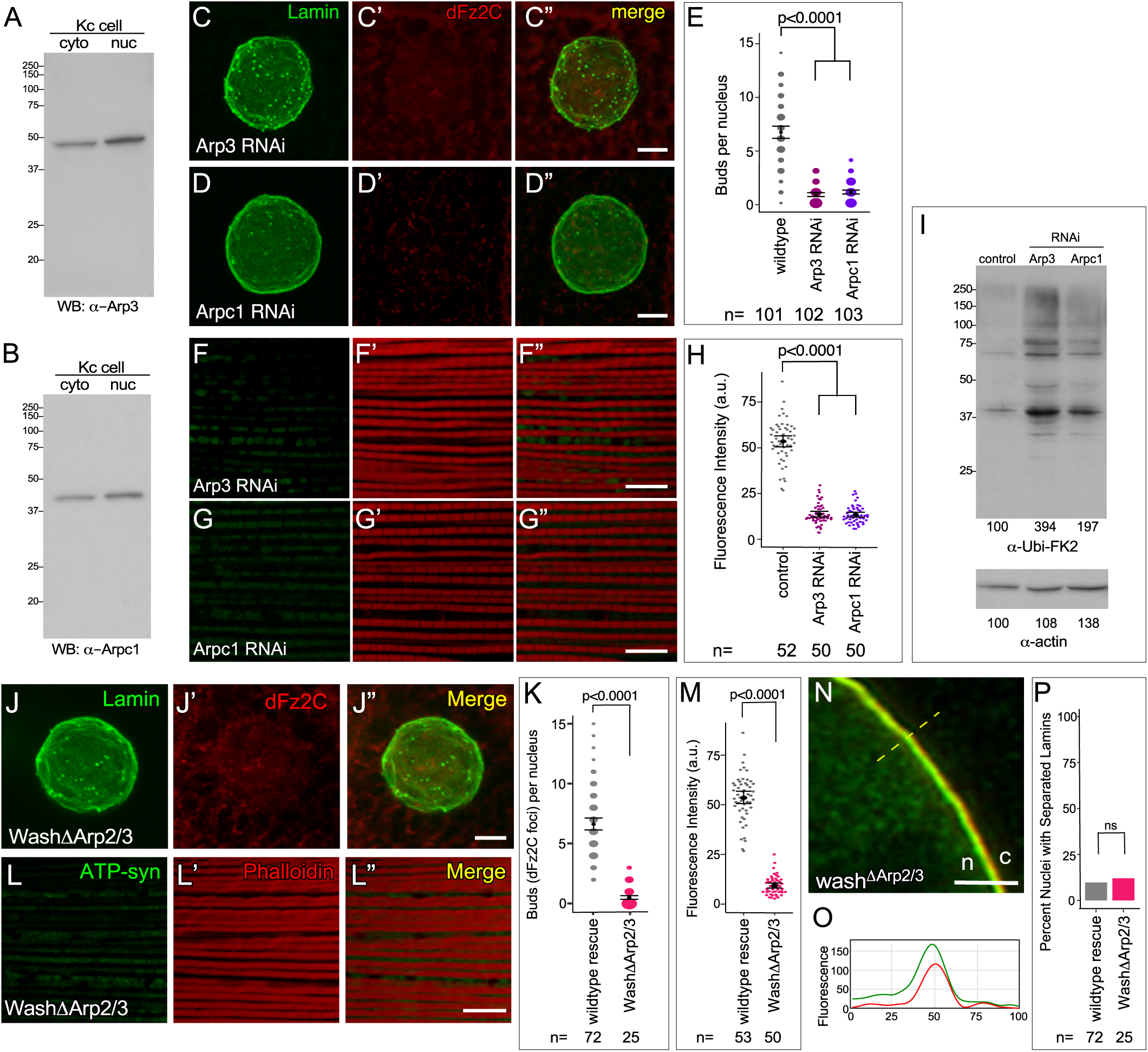
Arp2/3 activity is required for NE-budding. (A-B) Western blot of Kc cell cytoplasmic and nuclear extracts probed with antibodies to Arp2/3 subunit Arp3 (A) or Arpc1 (B). (C-D”) Confocal micrograph projections of Arp3 RNAi (C-C”) and Arpc1 RNAi (D-D”) salivary gland nuclei stained with Lamin Band dFz2C. (E) Quantification of NE-buds per nucleus in larval salivary glands. (F-G”) Confocal micrograph projections of adult IFM from Arp3 RNAi (F-F”) and Arpc1 RNAi (G-G”) flies aged 21 days stained with ATP-Syn α and Phalloidin. (H) Quantification of ATP-Syn α fluorescence intensity in adult IFM. (I) Western blot of adult IFM lysates from control, Arp3 RNAi and Arpc1 RNAi flies aged 21 days showing poly-ubiquitin aggregate protein levels, and actin loading control. (J-J”) Confocal micrograph projections of *wash*^ΔArp2/3^ larval salivary gland nucleus stained with Lamin B and dFz2C. (K) Quantification of NE-buds per nucleus in larval salivary glands. (L-L”) Confocal micrograph projections of adult IFM from *wash*^ΔArp2/3^ flies aged 21 days stained with ATP-Syn α and Phalloidin. (M) Quantification of ATP-Syn α fluorescence intensity in adult IFM. (N-O) Single slice super-resolution micrographs of larval salivary gland nucleus (N) and line plot of region indicated (O) from *wash*^ΔArp2/3^ showing Lamin B/Lamin C organization at the nuclear periphery. (P) Quantification of the percentage of salivary gland nuclei showing Lamin B/Lamin C separation. Two-tailed Fisher’s exact test (P); Two-tailed student’s t-test (H,M); Kruskal Wallis test (E,K); all p-values indicated. Scale bars: 5*μ*m in C-D”,J-J”; 1*μ*m in N; 10*μ*m in F-G”,L-L” (n=nucleus; c=cytoplasm).

As expected given their reduction of dFz2C foci/NE-buds, *Arp3* and *Arpc1* knockdowns also exhibit the phenotypes linked to disruption of NE-budding. IFM from 21 day old *Arp3* and *Arpc1* knockdown flies show a decrease in mitochondrial activity, as assayed using the activity dependent mitochondrial marker ATP-Synthetase α (Figure 7F-H). *Arp3* and *Arpc1* RNAi knockdown IFM’s show a 3.1 (n=50) and 3.9 (n=50) fold decrease in mitochondrial activity respectively (p<0.0001 in each case). Western blots from these tissues also show an increase in poly-ubiquitin aggregates, indicating mitochondrial damage (Figure 7I).

To confirm that Wash’s interaction with Arp2/3 is required for its role in NE-budding, we generated a point mutation in the conserved Tryptophan residue (W498S) that, when mutated in WASP, abolished binding of WASP to Arp2/3 (Marchand et al., 2001) (Figure S1I). This construct, harboring the point mutation in the context of the full-length Wash protein, designated wash^ΔArp2/3^, was examined for specificity using GST pulldown assays (Figure S1J). Wash^ΔArp2/3^ transgenics were generated in the same manner as for *wash^ΔSHRC^* and *wash^ΔΔLamB^*. Wash^ΔArp2/3^ functions as expected during oogenesis: it does not rescue the *wash* null ooplasmic streaming phenotype (Figure S1N-O).

Larval salivary gland nuclei from *wash^ΔArp2/3^* point mutants show significantly reduced NE-buds, with only 0.5±0.1 buds per nucleus (n=102, p<0.0001), and exhibited nuclear morphology indistinct from *wash^WT^* (Figure 7J-K). Consistent with a requirement for its actin nucleation activity during NE-budding, IFM from 21 day old *wash^ΔArp2/3^* point mutants show a decrease in mitochondrial activity, as assayed using the activity dependent mitochondrial marker ATP-Synthetase α (Figure 7L-M). IFM’s from *wash^ΔArp2/3^* show a 4.4 fold decrease in mitochondrial activity (n=50). Additionally, western blots of IFM lysates from *wash^ΔArp2/3^* show an increase in poly-ubiquitin aggregates, indicating mitochondrial damage (Figure 5P). As might be expected, *wash^ΔArp2/3^* does not exhibit separated Lamin B and Lamin C meshes (Figure 7N-P). Taken together, our data are consistent with a model in which Wash and the SHRC work upstream of the Arp2/3 complex to promote NE-budding, and that Wash functions independently of the SHRC and Arp2/3 to affect nuclear morphology.

## DISCUSSION

NE-budding is a newly appreciated pathway for nuclear export of large macromolecular machineries and/or cargoes, such as megaRNPs involved in the co-regulation of major developmental pathways or unwanted RNA/protein aggregates, that bypasses canonical nuclear export through nuclear pores. We previously showed that *Drosophila* Wash is present in the nucleus where, like in the cytoplasm, is likely involved in a number of different nuclear processes (Verboon et al., 2015b). Here, we show that all members of Wash’s four subunit regulatory complex (SHRC: CCDC53, Strumpellin, SWIP, FAM21) are also present within the nucleus and, along with Wash, are necessary for NE-budding. We show that Wash/SHRC act early in the NE-budding pathway and requires Wash’s actin nucleation activity achieved through its interaction with Arp2/3. Loss of Wash or any of its SHRC subunits leads to the loss of dFz2Cfoci/NE-buds, and mutants for these factors exhibit phenotypes associated with the two cellular processes shown to require NE-budding: aberrant synaptic “ghost” bouton formation leading to disrupted neuromuscular junction integrity, and mitochondrial degeneration associated with premature aging phenotypes (Li et al., 2016; Speese et al., 2012). While the spectrum of cellular/developmental processes that require this alternate nuclear egress mechanism is not yet known, SHRC components are linked to neurodegenerative disorders, including hereditary spastic paraplegias, Parkinson disease, amyotrophic lateral sclerosis (ALS), and Hermansky-Pudlak syndrome (Elliott et al., 2013; McGough et al., 2014; Ropers et al., 2011; Ryder et al., 2013; Song et al., 2018; Turk et al., 2017; Valdmanis et al., 2007; Vardarajan et al., 2012; Zavodszky et al., 2014a; Zavodszky et al., 2014b). As an increasing number of neurodegenerative diseases and myopathies have been associated with the accumulation of RNA-protein aggregates in the nucleus, NE-budding may be part of the endogenous cellular pathway for removing such aggregates/megaRNPs from the nucleus in normal cells (Laudermilch et al., 2016; Parchure et al., 2017; Ramaswami et al., 2013; Woulfe, 2008).

### Nuclear Buds or NPCs?

The parallels between NE-budding and the nuclear egress mechanisms used by herpesviruses, as well as the presence of similar endogenous perinuclear foci/buds in other plant and animal nuclei, has led to the suggestion that NE-budding might be a conserved endogenous cellular pathway for nuclear export independent of NPCs (Fradkin and Budnik, 2016; Panagaki et al., 2018; Parchure et al., 2017; Speese et al., 2012). However, INM-encapsulated electron-dense granules have been identified in yeast with similarities to the dFz2C foci/NE-buds observed in *Drosophila* muscle, salivary gland, and nurse cell nuclei (Parchure et al., 2017; Webster et al., 2014). These yeast granules contain NPC proteins, leading to the alternative suggestion that NE-budding is a means of removing defective NPCs. In addition, INM-tethered electron-dense granules have been described in Torsin-deficient HeLa cells, leading to the suggestion that NE-budding may also involve NPCs and/or NPC components (Laudermilch et al., 2016; Parchure et al., 2017). It has also been proposed that mis-regulation of NPC function could lead to an upregulation of NE-budding (Parchure et al., 2017). While the full relationship between NPCs and NE-buds is not yet known, one important difference is that the yeast and HeLa nuclear granules observed are much smaller (∼120nm) than dFz2C foci/NE-buds (∼500nm) (see Figure 1F-G). Our identification of Wash/SHRC, proteins with the capability of remodeling cortical cytoskeleton and/or membranes, which likely are part of the cellular machinery involved in physical aspects of NE-budding (as opposed to being a megaRNP components), lend support for NE-budding being an alternate endogenous nuclear exit pathway.

### The nuclear lamina and NE-budding

The handful of factors identified to date that play a role in NE-budding are mostly components of megaRNPs (Fradkin and Budnik, 2016; Parchure et al., 2017; Speese et al., 2012). The molecular and cellular machineries/mechanisms underpinning this process are largely unknown. NE-budding has been proposed to occur at sites along the INM where the nuclear lamina is modified by aPKC phosphorylation (Fradkin and Budnik, 2016; Parchure et al., 2017; Speese et al., 2012). Both A- and B- type lamins play a role in NE-budding and have been proposed to be the target of aPKC phosphorylation in the nuclear lamina, similar to PKC phosphorylation of lamins that precedes lamina disassembly in mitotic NE breakdown (Guttinger et al., 2009), apoptosis (Cross et al., 2000), or during viral capsid nuclear egress (Park and Baines, 2006). Viral NE-budding requires a virus-encoded Nuclear Egress Complex (NEC) consisting of two virally-encoded proteins (or their orthologs in different classes of herpesviruses), which have been implicated in the recruitment of viral and cellular kinases to the INM to modify the nuclear lamina (Bigalke and Heldwein, 2016; Mettenleiter et al., 2013; Zeev-Ben-Mordehai et al., 2015). Cellular counterparts for these virally-encoded NEC proteins have not yet been identified. It is also not yet known how this kinase activity is restricted to specific sites along the nuclear lamina or how those specific sites are selected.

We have previously shown that Wash interacts directly with Lamin B and that loss of nuclear Wash results in a wrinkled a nuclear morphology reminiscent of that observed in laminopathies (Verboon et al., 2015b). We reasoned that Wash’s disruption of the nuclear lamina may account for its NE-budding phenotypes. Consistent with this idea, we find Lamin B knockdown nuclei and nuclei from a *wash* point mutant that disrupts Wash’s interaction with Lamin B (wash^ΔΔLamB^) exhibit a wrinkled nuclear morphology, reduced dFz2C foci/NE-buds, and NE-budding associated phenotypes. Of note, their effects on NE-budding are not as strong as that of Wash or SHRC mutants.

Lamin A/C and Lamin B isoforms have been shown to form homotypic meshworks underlying the nuclear envelope that interact among themselves (in as yet unknown ways), and that are somehow linked to integral membrane proteins of the INM, the nuclear lamina/nucleoskeleton, and to the chromatin adjoining the INM (Shimi et al., 2015). Intriguingly, our data suggests that these lamin homotypic meshes are likely layered, rather than interwoven, and that Wash affects the anchoring of these different lamin homotypic meshes to each other and/or the INM. Absent dFz2C foci/NE-buds and strong NE-budding associated phenotypes are also observed in SHRC and Arp2/3 knockdown nuclei. However, these knockdowns do not exhibit the wrinkled nuclear morphology or separated lamin isoform meshes, suggesting that Wash can also affect NE-budding by a means independent of disrupted global nuclear lamina integrity. Interestingly, Wash’s functions with the SHRC and with Lamin B involve separate nuclear complexes. Taken together, our data suggest that loss of Wash affects NE-budding in two ways: making NE-budding inefficient by disrupting the nuclear lamina though its interaction with Lamin B, and disrupting the ability to form NE-buds through its interaction with its SHRC and Arp2/3 (see below). aPKC may play a similar, somewhat indirect, role in NE-budding by generally disrupting the nuclear lamina. Alternatively (or in addition to), aPKC may target Wash: WASH phosphorylation by Src kinases has been shown to be necessary for regulating NK cell cytotoxicity (Huang et al., 2016).

### A physical role for Wash in NE-budding

For bud formation/envelopment of a megaRNP or macromolecular cargo to occur the INM must interact with its underlying cortical nucleoskeleton (nuclear lamina) to allow the INM deformation/curvature necessary to form the physical NE-bud. Force must also be generated that allows the bud to extend into the perinuclear space, as well as for the pinching off (scission) of the nascent bud into the perinuclear space. In the cytoplasm, WAS family proteins are often involved in membrane-cortical cytoskeleton coupled processes, including both ‘inward’ membrane deformations (i.e., endocytosis, intracellular movement of pathogenic bacteria) and ‘outward’ membrane deformations (i.e., exocytosis, cell protrusions), required for signal/environment sensing and cell movement during normal development, as well as during pathological conditions such as metastasis (Burianek and Soderling, 2013; Campellone and Welch, 2010; Massaad et al., 2013; Rottner et al., 2010; Stradal et al., 2004; Takenawa and Suetsugu, 2007). WASH, in particular, has been implicated in endosome biogenesis and/or sorting in the cytoplasm, where it, along with its SHRC, drives Arp2/3-dependent actin assembly to influence endosome trafficking, remodels membrane, and facilitates membrane scission (Derivery et al., 2009; Duleh and Welch, 2010; Gomez and Billadeau, 2009; Rottner et al., 2010; Simonetti and Cullen, 2019). Thus, Wash encodes the biochemical properties needed to regulate the membrane deformation/curvature necessary to form the NE-bud and/or play a role in generating the forces necessary to pinch off the NE-bud from the INM. Consistent with Wash playing a role in the physical production of a NE-bud, we find that Wash acts prior to Torsin that is implicated in pinching off the NE-bud from the INM, and it requires its actin nucleation activity.

We have identified Wash and its SHRC as new players in the cellular machinery required for the newly described endogenous NE-budding pathway. Our data suggest that Wash is involved in two nuclear functions that can affect NE-budding. 1) Wash is required to maintain the organization of Lamin isoforms relative to each other and the INM through its direct physical interaction with Lamin B. This Wash activity is SHRC- and Arp2/3- independent and is likely a non-specific mechanism as global disruption of the nuclear lamina/nuclear envelope would indirectly affect many nuclear processes. 2) Wash is required for NE-bud formation. This Wash activity is SHRC- and Arp2/3- dependent. While focus of NE-budding research to date has centered on the composition of the megaRNPs and the spectrum of cellular/developmental processes requiring NE-budding, Wash/SHRC are likely involved in the physical aspects of NE-budding. Thus, Wash/SHRC provide a molecular entry into the physical machinery that underlies NE-budding. In the future, it will be exciting to further explore Wash’s roles in NE-budding, and to determine how it functions to get macromolecular complexes through the INM, and how closely these nuclear roles parallel those in the cytoplasm.

## MATERIALS AND METHODS

### KEY RESOURCES TABLE

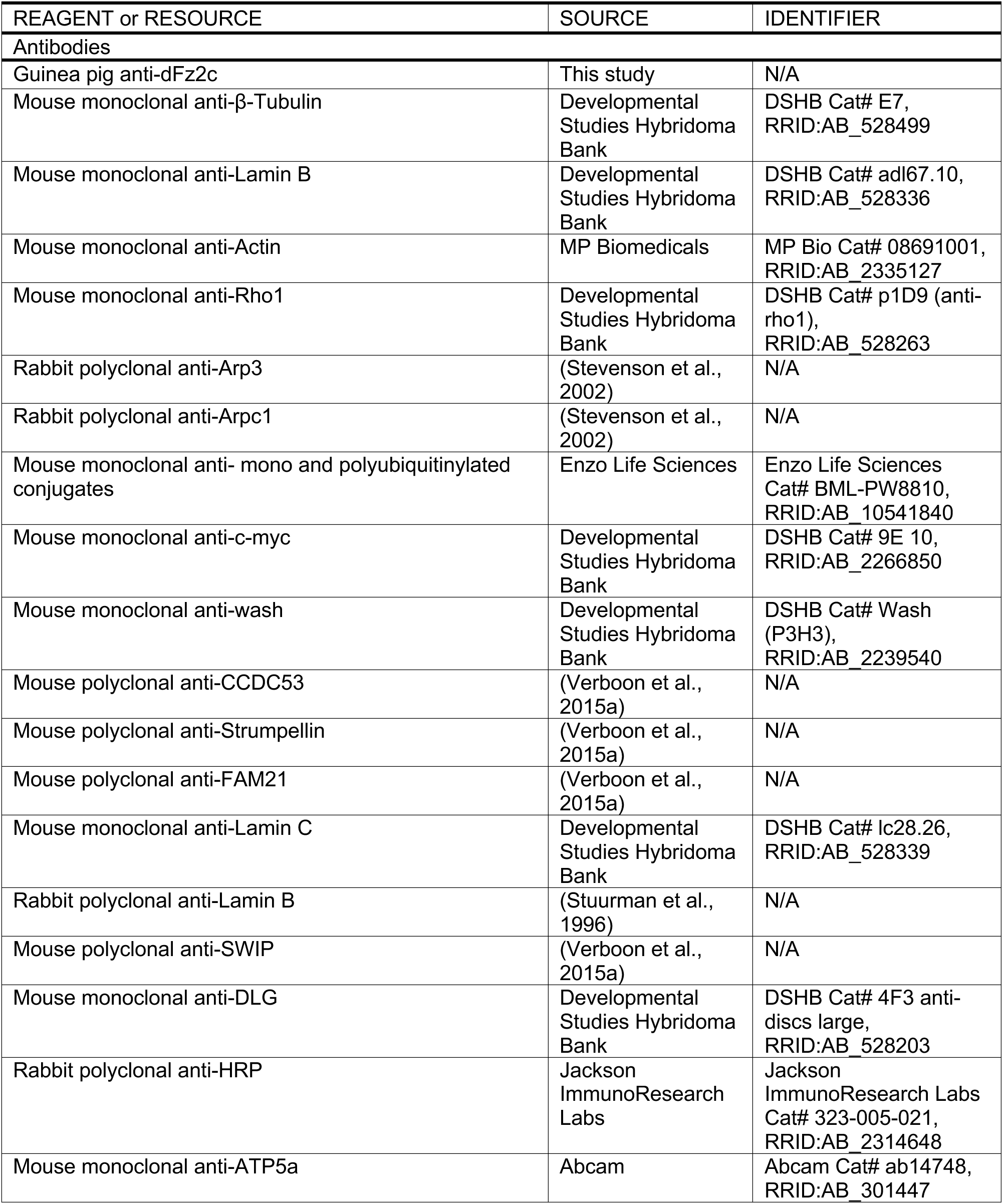
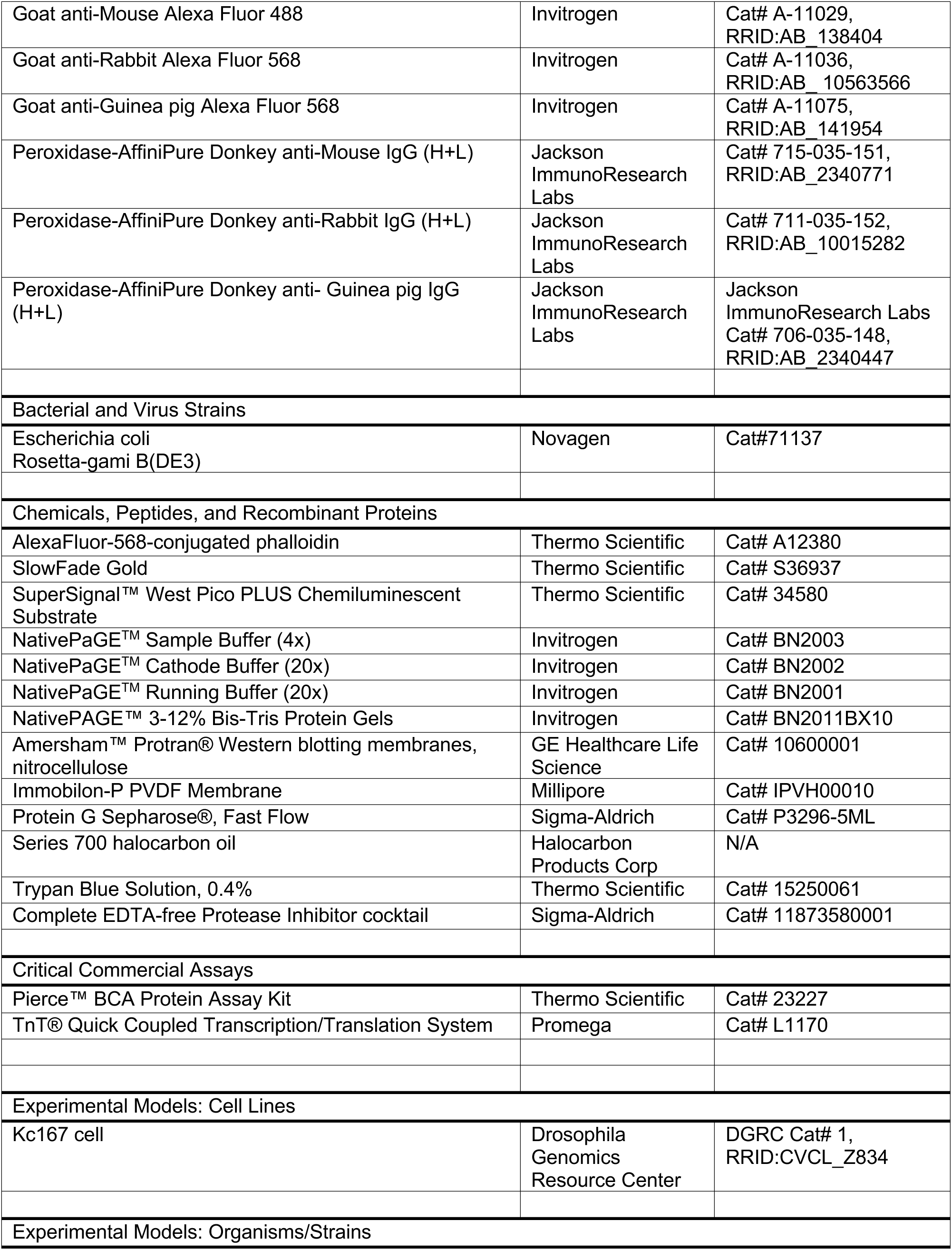
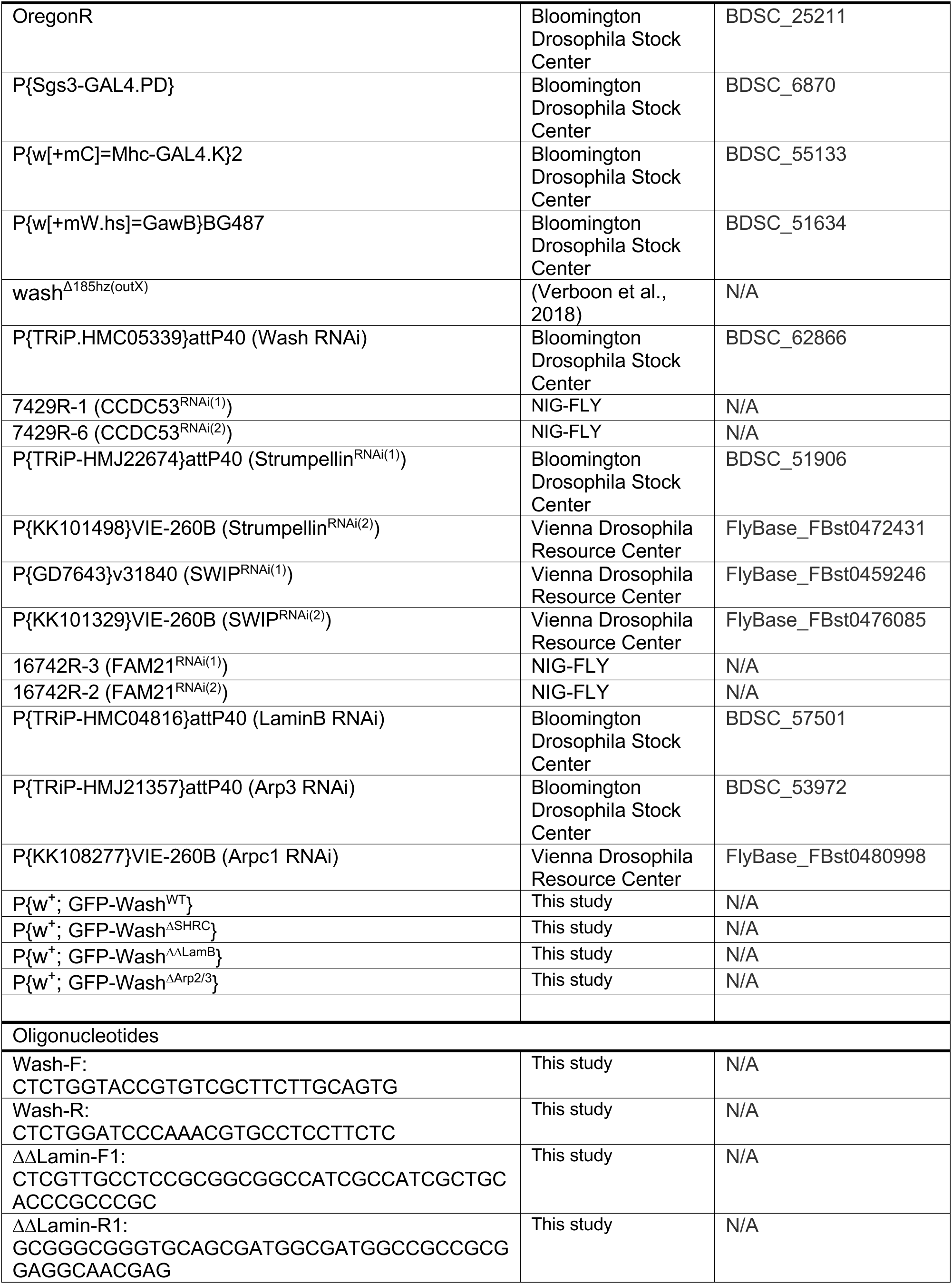
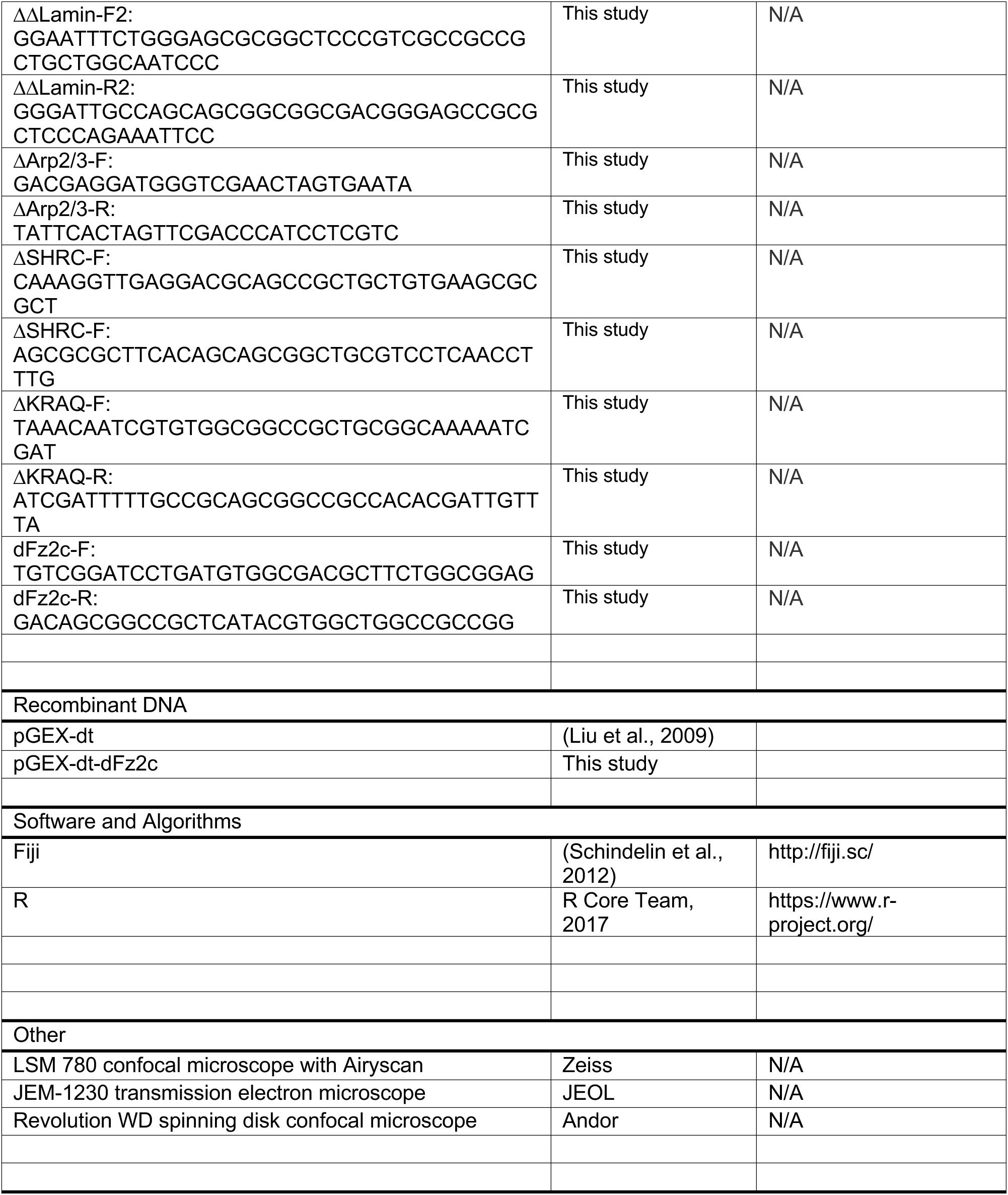

### Fly stocks and genetics

Flies were cultured and crossed at 25°C on yeast-cornmeal-molasses-malt extract medium. Flies used in this study are listed in the Key Resources Table. All fly stocks were treated with tetracycline and then tested by PCR to ensure that they did not harbor Wolbachia. RNAi knockdowns were driven in the salivary glands by the GAL4-UAS system using the P{Sgs3-GAL4.PD} driver (Bloomington *Drosophila* Stock Center, stock #6870). RNAi knockdowns were driven in the indirect flight muscle by the GAL4-UAS system using the P{w[+mC]=Mhc-GAL4.K}2 driver (Bloomington *Drosophila* Stock Center, stock #55133). RNAi knockdowns were driven in the larval body wall muscle by the GAL4-UAS system using the P{w[+mW.hs]=GawB}BG487 driver (Bloomington *Drosophila* Stock Center, stock #51634). The *wash^Δ185^* deletion allele was kept as a continuously outcrossed stock (Verboon et al., 2018).

### Construction of *Wash* point mutant transgenic lines

The residues required for Wash’s interaction with Lamin and CCDC53 were mapped by successive GST pulldown assays using fragments of Wash protein, followed by specific point or substitution mutations in the context of the full-length Wash protein (as detailed in Figures 6A and 7J). Point and/or substitution mutations were confirmed by sequencing.

A 2.9 kb genomic fragment encompassing the entire Wash gene was PCR’d, then subcloned into the Casper 4 transformation vector by adding KpnI (5’) and BamHI (3’) restriction sites. GFP was inserted N-terminal to the Wash ATG by PCR (GFP-Wash^WT^). The Wash portion of this construct was swapped with the point and/or substitution mutations described above to generate GFP-Wash^ΔSHRC^, GFP-Wash^ΔΔLamB^, and GFP-Wash^ΔArp2/3^.

These constructs were used to make germline transformants as previously described (Spradling, 1986). Transgenic lines that mapped to Chromosome 2 and that had non-lethal insertions were kept. The resulting transgenic lines (P{*w^+^*; *GFP-Wash^WT^*}, P{*w^+^*; *GFP-Wash^ΔSHRC^*}, P{*w^+^*; *GFP-Wash^ΔΔLamB^*}, and P{*w^+^*; *GFP-Wash ^ΔArp2/3^*}) were recombined onto the *wash^Δ185^* null chromosome to assess the contribution of the particular Wash transgene. The resulting recombinants (*wash^Δ185^* P{*w^+^*; *GFP-Wash^WT^*}) are essentially gene replacements, as *wash* activity is only provided by the transgene. These transgenes do not rely on over-expression, but rather on the spatial and temporal expression driven by the endogenous *wash* promoter itself. We analyzed a minimum of three lines per construct and have checked all lines to confirm that the levels and spatial distribution of their expression is indistinguishable from wildtype. The wildtype version of this transgene (*wash^Δ185^* P{ *w^+^*; *GFP-Wash^WT^*}) rescues the phenotypes associated with the outcrossed *wash^Δ185^* mutation (Liu et al., 2009; Verboon et al., 2018; Verboon et al., 2015a; Verboon et al., 2015b).

### Antibody production and characterization

Guinea pig antibodies were raised against bacterially purified Flag-tagged dFz2c protein at Pocono Rabbit Farm & Laboratory (Canadensis, PA) using their standard protocols. Protein was purified as described previously (Rosales-Nieves et al., 2006). Western blotting was used to confirm antibody specificity using dFz2 purified protein, Kc cell, ovary, and S2 whole cell extracts. Antibody generation to Wash and SHRC subunits were described previously (Rodriguez-Mesa et al., 2012; Verboon et al., 2015a).

### Lysate preparation

*Drosophila* cytoplasmic and nuclear extracts were made from Kc167 cells. Briefly, cells were grown to confluence in 500 ml spinflasks, pelleted for 5 min at 1500 rpm, resuspended in 100 ml cold 1× PBS and re-pelleted for 5 min at 1500 rpm. Cell pellets were flash frozen in liquid nitrogen. Cells were resuspended in Sucrose Buffer (0.32M Sucrose, 3mM CaCl_2_, 2mM MgAc, 0.1mM EDTA, 1.5% NP40) with 2× protease inhibitors [Complete Protease Inhibitor (EDTA free; Sigma, St. Louis, MO), 2mM PMSF, and 1mM Na_3_VO_4_] and 2× phosphatase inhibitors [PhosSTOP; Sigma, St. Louis, MO] at 100 μL per 1 million cells and incubated on ice for 30 min. Lysate was dounce homogenized 10× on ice. Lysate was then centrifuged 10 min at 3750 rpm at 4°C, nuclei formed a pellet and supernatant was cytoplasmic extract. Lipids were removed from the top of cytoplasmic extract using a sterile swab, then cytoplasmic fraction was removed and centrifuged for 10 minutes at 4000 rpm at 4°C. Cytoplasmic supernatant was removed and 1/10^th^ of the supernatant volume of 11× RIPA was added. Cytoplasmic extract was aliquoted and flash frozen. Nuclear pellet was resuspended with Sucrose Buffer with protease/phosphatase inhibitors and NP40 and re-dounced. Nuclear lysate was then centrifuged for 10 minutes at 4000 rpm at 4°C, and supernatant was discarded. Nuclear pellet was resuspended with Sucrose buffer without NP40 and centrifuged for 20 minutes at 4000 rpm at 4°C. Supernatant was discarded and nuclear pellet was resuspended in 2.5mL of (20mM HEPES pH 7.9, 0.5mM EDTA, 100mM KCl, and 10% Glycerol) per liter of cells used. DNA was degraded using MNase in a 37°C water bath for 10 minutes. 20μL of 500mM EDTA per 500μL lysate was added and incubated on ice for 5 minutes. Lysate was then nutated for 2 hours at 4°C. Lysate was sonicated using a Sonic Dismembrator (Model 60; Fisher Scientific) at setting 3.5 with 10 s per pulse for 15 minutes. Lysate was clarified with a 15 min centrifugation at 15,000 rpm, aliquoted and flash frozen.

Adult indirect flight muscle (IFM) lysate was made from 21 day old flies in RIPA buffer (HEPES 50mM pH 7.5, NaCl 150 nM, 1% NP40, 0.5% Deoxycholate sodium salt, EDTA 5mM). IFM was dissected from flies in cold PBS. PBS was removed and 200μL of 1× RIPA buffer with 2× protease inhibitors [Complete EDTA free Protease inhibitor; Sigma, St. Louis, MO] per 20 IFMs was added. Lysates were homogenized with an Eppendorf microcentrifuge homogenizing pestle on ice. Lysates were then sonicated with a probe sonicator on setting 3 for three 10-second pulses and centrifuged at 15,000 rpm for 30 minutes at 4°C. Supernatant was removed and MgCl_2_ was added to 2mM and 8μL of Benzonase (Millipore) was added per 200μL of lysate. Lysates were nutated at room temperature for 15 minutes. Lysates were centrifuged at 15,000 rpm for 30 minutes at 4°C. Supernatant was aliquoted and flash frozen.

### Western blotting

Cytoplasmic and nuclear purity of lysates was assayed using (β-Tubulin E7, 1:2000, Developmental Studies Hybridoma Bank) and (Lamin B monoclonal 67.10, 1:1000, Developmental Studies Hybridoma Bank). Lysate samples were normalized to a loading control (Actin Clone C4, 1:2500; MP Biochemicals) and then blotted according to standard procedures. The following antibodies were used: anti-Rho1 monoclonal (P1D9, 1:50, Developmental Studies Hybridoma Bank), anti-Arp3 (1:500, (Stevenson et al., 2002)), and anti-Arpc1 (1:500, (Stevenson et al., 2002)), anti-mono and polyubiquitinylated conjugates (FK2, 1:1000, Enzo Life Sciences, East Farmingdale, NY). Quantification of actin loading controls and Ubiquitin-FK2 expression was done using ImageJ (NIH).

### Immunoprecipitation

Nuclear lysate was incubated with primary antibody overnight at 4°C. Protein G sepharose (20 μL) was then added in 0.5 ml Carol Buffer (50 mM HEPES pH 7.9, 250 mM NaCl, 10 mM EDTA, 1 mM DTT, 10% glycerol, 0.1% Triton X-100) + 0.5 mg/ml bovine serum albumin (BSA) + protease inhibitors (Complete EDTA-free Protease Inhibitor cocktail; Sigma, St. Louis, MO) and the reaction allowed to proceed for 2 hours at 4°C. The beads were washed 1× with Carol Buffer + BSA and 2× with Carol Buffer alone. Analysis was conducted using SDS-PAGE followed by Western blots. Antibodies used for immunoprecipitations are as follows: anti-9e10 (Developmental Studies Hybridoma Bank), anti-Wash monoclonal (P3H3, (Rodriguez-Mesa et al., 2012)), anti-CCDC53 (Verboon et al., 2015a), anti-Strumpellin (Verboon et al., 2015a), anti-FAM21 (Verboon et al., 2015a), anti-Lamin B monoclonal (AD67.10, Developmental Studies Hybridoma Bank), anti-Lamin C monoclonal (LC28.26, Developmental Studies Hybridoma Bank). Antibodies used for the IP western blots are as follows: mouse anti-CCDC53 polyclonal (1:1000), mouse anti-Strumpellin polyclonal (1:400), mouse anti-FAM21 polyclonal (1:400), and anti-Lamin B monoclonal 67.10 (1:200).

### GST pulldown assays and Blue Native PAGE

GST pulldown assays were performed as previously described (Magie and Parkhurst, 2005; Magie et al., 2002; Rosales-Nieves et al., 2006). Blue Native Page was performed using a Novex Native PAGE Bis-Tris Gel System (Invitrogen) following manufacturer protocols. Briefly, *Drosophila* Kc cell nuclear extract was resuspended in 1× NativePAGE Sample Buffer (Invitrogen) with 1% digitonin and protease inhibitors, and incubated for 15 min on ice. Samples were centrifuged at 16,200 × *g* for 30 min at 4°C, and supernatant was resuspended with G250 sample additive and Native PAGE Sample Buffer. These prepared samples were loaded on 3–12% Bis-Tris Native PAGE gels and electrophoresed using 1×Native PAGE Running buffer system (Invitrogen). The cathode buffer included 1×Cathode Buffer Additive (Invitrogen). Native Mark Protein standard (Invitrogen) was used as the molecular weight marker. Protein concentrations of adult fly mitochondrial preps were determined with a BCA Protein Assay Kit (Thermo Scientific, USA) following manufacturer instructions. The following antibodies were used: mouse anti-Wash monoclonal (P3H3, 1:2, (Rodriguez-Mesa et al., 2012)), mouse anti-CCDC53 polyclonal (1:400, (Verboon et al., 2015a)), mouse anti-Strumpellin polyclonal (1:400, (Verboon et al., 2015a)), mouse anti-FAM21 polyclonal (1:400, (Verboon et al., 2015a)), rabbit anti-Lamin B polyclonal (L6, 1:2500, (Stuurman et al., 1996)).

### Electron Microscopy

*Drosophila* third instar larva salivary glands were processed for electron microscopy essentially as previously described (Pitt et al., 2000). Glands were dissected in 1x PBS then placed directly in fixative solutions [2.2% glutaraldehyde, 0.9% paraformaldehyde, 0.05 M cacodylate (pH 7.4), 0.09M sucrose, 0.9 mM MgCl2]. Glands were fixed for 2.5 h at room temperature, followed by several rinses with 0.09 M sucrose, 0.05 M cacodylate (pH7.4). Glands were post-fixed in 1% osmium, 0.8% potassium ferricyanide, 0.1 M cacodylate (pH 7.2) for 45 min at 4°C, followed by several rinses in 0.05 M cacodylate (pH 7.0). Glands were then treated with 0.2% tannic acid in 0.05 M cacodylate (pH 7.0) for 15 min at room temperature, followed by several rinses in dH_2_O. Glands were placed in 1% uranyl acetate in 0.1 M sodium acetate (pH 5.2) for 1h at room temperature, and rinsed three times with 0.1 M sodium acetate (pH 5.2), followed by three rinses with dH_2_O. Specimens were dehydrated in a graded acetone series, embedded in Epon, and sectioned following standard procedures. Grids were viewed with a JEOL JEM-1230 transmission electron microscope and photographed with a Gatan UltraScan 1000 CCD camera.

### Immunostaining of larval salivary glands

Salivary glands were dissected, fixed, stained, and mounted as previously described (Verboon et al., 2015a). Primary antibodies were added at the following concentrations: mouse anti-Wash monoclonal (P3H3, 1:200, (Rodriguez-Mesa et al., 2012)), mouse anti-CCDC53 polyclonal (1:300, (Verboon et al., 2015a)), mouse anti-Strumpellin polyclonal (1:300, (Verboon et al., 2015a)), mouse anti-SWIP polyclonal (1:300, (Verboon et al., 2015a)), mouse anti-FAM21 polyclonal (1:300, (Verboon et al., 2015a)), mouse anti-Lamin B monoclonal (AD67.10 1:200, Developmental Studies Hybridoma Bank), mouse anti-Lamin C monoclonal (LC 28.26, 1:200, Developmental Studies Hybridoma Bank), and guinea pig anti-dFz2C (1:2500, this study).

### Immunostaining of larval body wall muscle

Flies were transferred daily and wandering 3rd instar larvae were collected and subsequently fileted in cold PBS. Body wall muscle filets were fixed for 10 minutes. Fix is: 16.7 mM KPO4 pH6.8, 75 mM KCl, 25 mM NaCl, 3.3 mM MgCl2, 6% formaldehyde. After three washes with PTW (1× PBS, 0.1% Tween-20), larval filets were permeabilized in 1× PBS plus 1% Triton X-100 for 2 hours at room temperature, then blocked using PAT (1× PBS, 0.1% Tween-20, 1% BSA, 0.05% azide) for 2 hours at 4°C. Antibodies were added at the following concentrations: mouse anti-DLG (1:50), rabbit anti-HRP (1:1400). The larval filets were incubated 48 hours at 4°C. Primary antibody was then removed and filets were washed three times with XNS (1× PBS, 0.1% Tween-20, 0.1% BSA, 2% normal goat serum) for 30 min each. Alexa conjugated secondary antibodies (Invitrogen, Carlsbad, CA) diluted in PbT (1:1000) were then added and the filets were incubated overnight at 4°C. Larval filets were washed 10 times with PTW at room temperature for 10 minutes each and were mounted on slides in SlowFade Gold media (Invitrogen, Carlsbad, CA) and visualized using a Zeiss confocal microscope as described below. Total number of boutons was quantified in third instar larval preparations double labeled with antibodies to HRP and DLG, at segments A2-A3 muscles 6-7. The number of ghost boutons was assessed by counting HRP positive, DLG negative boutons.

### Actin visualization and immunostaining of indirect flight muscle

UAS controlled RNAi expressing flies were crossed to MHC-GAL4 driver flies and female RNAi/MHC-GAL4 trans-heterozygous flies were collected and aged 21 days at 25°C. Fly thoraxes were dissected and cut along the ventral side in cold PBS. Thoraxes were fixed using 1:6 fix/heptane for 15 minutes. Fix is: 16.7 mM KPO4 pH6.8, 75 mM KCl, 25 mM NaCl, 3.3 mM MgCl2, 6% formaldehyde. After three washes with PTW (1× PBS, 0.1% Tween-20) thoraxes were then cut along the dorsal side resulting in two halves and fixed again for 10 minutes using 1:6 fix/heptane for 15 minutes. After three washes with PTW (1× PBS, 0.1% Tween-20), thoraxes were permeabilized in 1× PBS plus 1% Triton X-100 for 2 hours at room temperature, then blocked using PAT (1× PBS, 0.1% Tween-20, 1% BSA, 0.05% azide) for 2 hours at 4°C. Antibodies were added at the following concentrations: mouse anti-ATP-Synthase α (15H4C4 1:100, Abcam, Cambridge, UK), and mouse anti-mono and polyubiquitinylated conjugates (FK2 1:200, Enzo Life Sciences, East Farmingdale, NY). The thoraxes were incubated 48 hours at 4°C. Primary antibody was then removed and thoraxes were washed three times with XNS (1× PBS, 0.1% Tween-20, 0.1% BSA, 2% normal goat serum) for 30 min each. Alexa conjugated secondary antibodies (Invitrogen, Carlsbad, CA) diluted in PbT (1:1000) and Alexa conjugated Phalloidin (1:50) were then added and the thoraxes were incubated overnight at 4°C. Thoraxes were washed 10 times with PTW at room temperature for 10 minutes each and were mounted on slides in SlowFade Gold media (Invitrogen, Carlsbad, CA) and visualized using a Zeiss confocal microscope as described below. To quantify ATP-Synthase α expression, we measured 512×512 pixel regions of the IFM and measured total fluorescence using ImageJ (NIH). Fluorescence measurements from separate experiments were normalized to the control genotype.

### Confocal and Super-resolution microscopy

Images of fixed tissues were acquired using a Zeiss LSM 780 spectral confocal microscope (Carl Zeiss Microscopy GmbH, Jena, Germany) fitted with a Zeiss 40×/1.0 oil Plan-Apochromat objective and a Zeiss 63×/1.4 oil Plan-Apochromat objective. FITC (Alexa Fluor 488) fluorescence was excited with the 488 nm line of an Argon laser, and detection was between 498 and 560 nm. Red (Alexa Fluor 568) fluorescence was excited with the 561 nm line of a DPSS laser and detection was between 570 and 670 nm. Pinhole was set to 1.0 Airy Units. Confocal sections were acquired at 0.2-1.0 μm spacing. Super-resolution images were acquired using an Airyscan detector in Super Resolution mode and captured confocal images were then processed using the Airyscan Processing feature on the Zen software provided by the manufacturer (Carl Zeiss Microscopy GmbH, Jena, Germany).

### Live image acquisition

To obtain live time-lapse images of oocytes, female flies were first fattened on yeast for 2 days. Females were then injected in the abdomen with 0.4% Trypan Blue (ThermoFisher Scientific, Waltham, MA) diluted 1:5 in PBS, and allowed to sit for 1-2 hours. Ovaries were dissected into individual egg chambers in halocarbon 700 oil (Halocarbon Products, River Edge, NJ) on a cover slip. Images were acquired on a Revolution WD systems (Andor Technology Ltd., Concord, MA) mounted on a Leica DMi8 (Leica Microsystems Inc., Buffalo Grove, IL) with a 63x/1.4 NA objective lens with a 2x coupler and controlled by MetaMorph software. Images were acquired with 561 nm excitation using an Andor iXon Ultra 888 EMCCD camera (Andor Technology Ltd., Concord, MA, USA). Time-lapse images were obtained by taking one single frame acquisition every 10 seconds for either 5 or 30 min.

### Statistical analysis

All statistical analyses were performed using R-3.6.1. For count data (# nuclear buds, # ubi puncta), gene knockdowns and point mutations were compared to the appropriate control, and statistical significance was calculated using a Kruskal Wallis test for independence. For frequency of observation (ghost vs. mature boutons, swirling vs. not swirling, lamin separated vs overlapping) a two-tailed Fisher’s Exact test was used. For all others, a two-tailed student’s t-test was used to test for significance between conditions. For plots, the mean + 95% CI was calculated using the Hmisc-4.2-0 package using 1000 bootstrap resamples. p < 0.01 was considered significant.

## ACKNOWLEDGMENTS

We thank Jim Priess, Bobbie Schneider, and Steve MacFarlane for help with EM, and Helen McNeill, Yonit Tsatskis, Bina Sugumar, and Clara Prentiss for help with BN-PAGE. We thank Vivian Budnik, Lynn Cooley, Paul Fisher, Bill Theurkauf, the Bloomington Stock Center, Kyoto Stock Center, Harvard Transgenic RNAi Project, Drosophila Genomics Resource Center, and Developmental Studies Hybridoma Bank for antibodies, DNAs, flies, and other reagents used in this study. This work was supported, in part, by NIH grant R01 AG023779 (to SMP) and by NCI Cancer Center Support Grant P30 CA015704 (Pilot project and Shared Resources).

## AUTHOR CONTRIBUTIONS

JMV and SMP designed the study and wrote the manuscript. All authors performed experiments, analyzed data, participated in discussion of the results, and commented on the manuscript.

---

**Figure S1.**
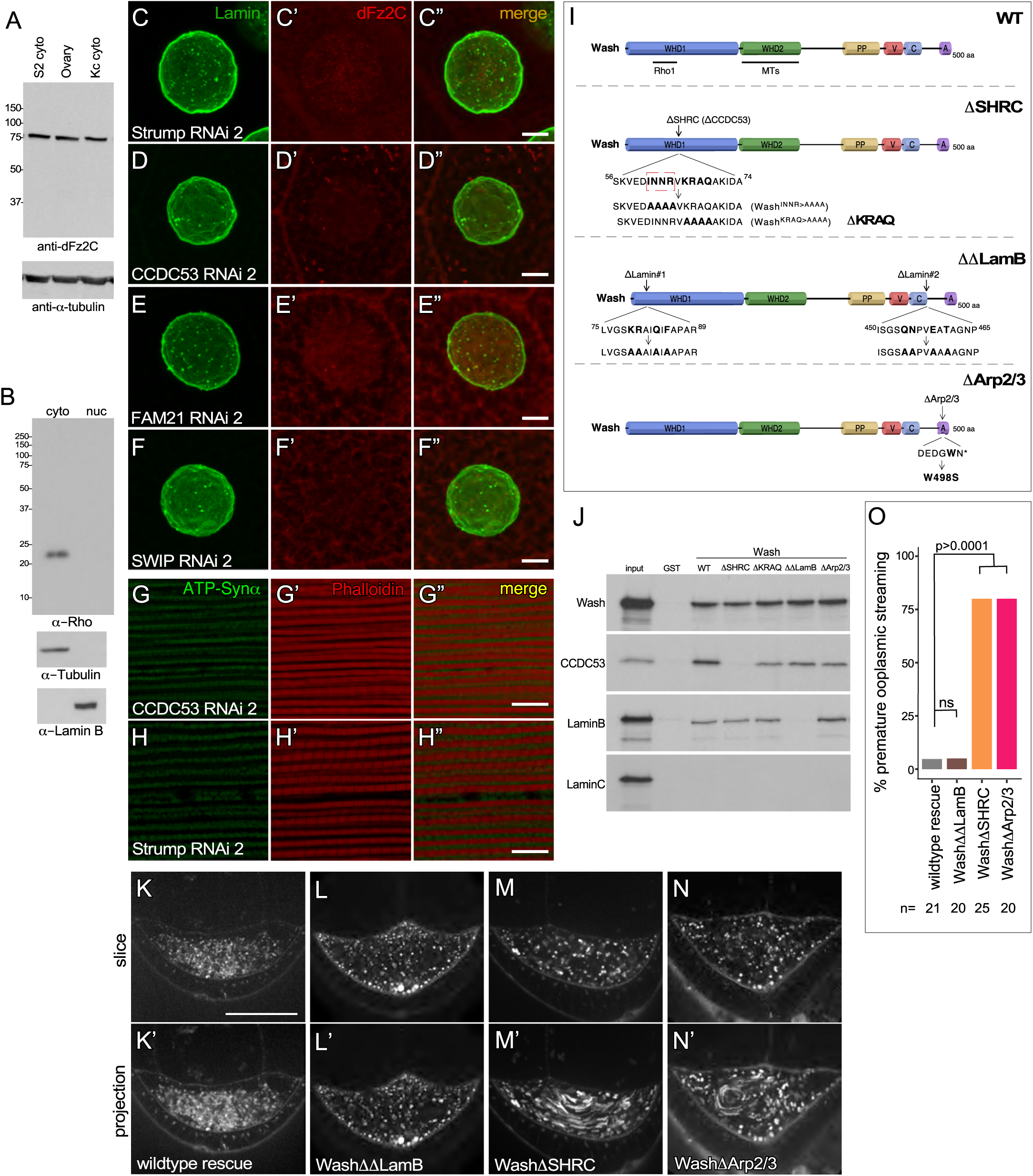
dFz2C antibody specificity, SHRC mutants lack buds and show NE-budding phenotypes, and Wash point mutant characterization. (A) Western blot of S2 cell cytoplasmic, ovary whole cell, and Kc cell cytoplasmic extracts probed with antibodies to dFz2C to show antibody specificity. Loading control is α-Tubulin. (B) Western blot of Kc cell cytoplasmic and nuclear extracts probed with antibodies to Rho1. Blots probed with antibodies to Tubulin and Lamin show cytoplasm and nuclear purity of extracts. (C-F”) Confocal micrograph projections of CCDC53 RNAi 2 (C-C”), Strumpellin RNAi 2 (D-D”), SWIP RNAi 2 (E-E”), and FAM21 RNAi 2 (F-F”) larval salivary gland nuclei stained with Lamin Band dFz2C. (G-H”) Confocal micrograph projections of adult IFM from CCDC53 RNAi 2 (G-G”) and Strumpellin RNAi 2 (H-H”) flies aged 21 days stained with activity dependent mitochondrial marker ATP-Syn α and Phalloidin. (I) Schematic diagram of the Wash wildtype (WT) rescue and point/substitution mutation constructs used to generate the transgenic lines indicating the position and specific substitution mutations for each construct (not drawn to scale). (J) GST pulldown experiments demonstrating the regions of Wash required for binding to Wash, CCDC53 and Lamin B, as well as the lack of direct binding of Wash by Lamin C. ^35^S-labeled *in vitro* translated Wash, CCDC53, Lamin B and Lamin C were tested with bacterially purified GST Wash (wildtype or point mutation containing) proteins as indicated. 10% input is shown. (K-N’) Single time-point (K-N) and 30 time-point projections (K’-N’) of live time-lapse movies of stage 7 oocytes in *wash*^WT^ (K-K’), *wash*^ΔΔLamB^, (L-L’), *wash*^ΔSHRC^ (M-M’), and *wash*^ΔArp2/3^ (N-N’). (O) Quantification of the percentage of stage 7 oocytes exhibiting premature ooplasmic streaming (N= for each genotype indicated on graph). Two-tailed Fishers exact test; p-values indicated. Scale bars: 5μm in C-F”; 10μm in G-H”; 50μm in K-N’.

